# A distinct class of bursting neurons with strong gamma synchronization and stimulus selectivity in monkey V1

**DOI:** 10.1101/583955

**Authors:** Irene Onorato, Sergio Neuenschwander, Jennifer Hoy, Bruss Lima, Katia-Simone Rocha, Ana Clara Broggini, Cem Uran, Georgios Spyropoulos, Thilo Womelsdorf, Pascal Fries, Cristopher Niell, Wolf Singer, Martin Vinck

**Affiliations:** Ernst Strüngmann Institute (ESI) for Neuroscience in Cooperation with Max Planck Society, Frankfurt, Germany; Max Planck Institute for Brain Research, Frankfurt, Germany; Frankfurt Institute for Advanced Studies, Frankfurt, Germany; Brain Institute, Federal University of Rio Grande do Norte, Natal, Brazil; Vanderbilt University, Nashville, USA; Institute of Neuroscience and Department of Biology, University of Oregon, Eugene

## Abstract

Cortical computation depends on interactions between excitatory and inhibitory neurons. The contributions of distinct neuron-types to sensory processing and network synchronization in primate visual-cortex remain largely undetermined. We show that in awake monkey V1, there exists a distinct cell-type (≈30% of neurons) that has narrow-waveform action-potentials, high spontaneous discharge-rates, and fires in high-frequency bursts. These neurons are more stimulus-selective and phase-locked to gamma (30-80Hz) oscillations as compared to other neuron types. Unlike the other neuron-types, their gamma phase-locking is highly predictive of their orientation tuning. We find evidence for strong rhythmic inhibition in these neurons, suggesting that they interact with interneurons to act as excitatory pacemakers for the V1 gamma rhythm. These neurons have not been observed in other primate cortical areas and we find that they are not present in rodent V1. However, they resemble the excitatory “chattering” neurons previously identified by intracellular recordings in cat V1. Given its properties, this neuron type should be pivotal for the encoding and transmission of V1 stimulus information.

## Introduction

Cortical tissue contains different types of neurons, which can be distinguished based on molecular, electrophysiological and histological markers (Batista-Brito et al., 2018; Moore et al., 2010; Rudy et al., 2011; Gentet, 2012; Markram et al., 2004). Interactions between the two main classes, excitatory (E) and inhibitory (I) neurons, govern sensory responses and the emergence of network oscillations and synchrony (Isaacson and Scanziani, 2011; Wilson et al., 2018; Vinck et al., 2013b; Buzsáki and Wang, 2012; Kopell et al., 2000; Wang, 2010; Haider et al., 2010; Haider and McCormick, 2009). However, the nature of these E-I interactions in primate model systems remains largely undetermined.

The primary visual cortex is one of the most widely studied model systems of the nervous system. The response properties of V1 neurons, for example orientation tuning, have been extensively described in different species. Furthermore, many visual stimuli give rise to prominent fast “gamma” oscillations (30-80Hz) in area V1 (Gray et al., 1989; Jagadeesh et al., 1992; Gray et al., 1990). The oscillatory synchronization of V1 spiking-activity may determine how stimulus information is transmitted to higher brain areas (Fries, 2015; Womelsdorf et al., 2012; Salinas and Sejnowski, 2000; Singer and Gray, 1995; Abeles, 1982; Buzsáki, 2006). E-I interactions are thought to make major contributions to both V1 orientation tuning and gamma oscillations (Shapley et al., 2003; Buzsáki and Wang, 2012; Wang, 2010). Recent studies in rodents have began to genetically target distinct cell-types to study their precise contributions to sensory tuning and network oscillations. These studies support a role for inhibition in shaping orientation- and direction-selectivity of V1 neurons (Kerlin et al., 2010; Znamenskiy et al., 2018; Perrenoud et al., 2016). Furthermore, they have implied specific classes of inhibitory interneurons in the generation of V1 gamma oscillations in mice (Veit et al., 2017; Chen et al., 2017; Senzai et al., 2019; Perrenoud et al., 2016). Yet, rodents and primates differ greatly in the functional and anatomical organization of their cortices (Ohki and Reid, 2007), and in their cognitive capacity. Further, monkey V1 shows gamma oscillations that are very strong as compared to other model systems like monkey V4 and mouse V1 (Vinck and Bosman, 2016), suggesting that the nature of E-I interactions might be fundamentally different in monkey V1. It is thus critical to study the contributions of different cell types to sensory processing and synchronization in primate V1. In primate cortex there are only limited possibilities to target cell types based on molecular markers. Primate studies have therefore analyzed action potential waveforms and firing statistics to distinguish between fast-spiking interneurons and excitatory cells, which generally yields high overlap with genetic and morphological markers of cell classes (McCormick et al., 1985; Senzai et al., 2019; Miri et al., 2018; Gentet et al., 2012; Cardin et al., 2009; Vinck et al., 2013a; Ardid et al., 2015; Mitchell et al., 2007; Vinck et al., 2016; Perrenoud et al., 2016; Hasenstaub et al., 2005; Nowak et al., 2003; Trainito et al., 2019). Furthermore, the differentiation between distinct excitatory cell types depends at present primarily on firing characteristics (Nowak et al., 2003; Bartho et al., 2004).

In the present study, we recorded spikes and local field potential (LFP) activity from awake macaque V1. We analyzed firing statistics and action-potential waveforms to distinguish between different cell classes. Fast-spiking interneurons and excitatory cells are commonly distinguished by their narrow- and broad-waveform spikes, respectively (Senzai et al., 2019; McCormick et al., 1985; Csicsvari et al., 1999; Perrenoud et al., 2016). Yet, we observed that the percentage of narrow-waveform neurons in macaque V1 largely exceeds the known percentage of GABAergic interneurons (Hendry et al., 1987). We show that narrow-waveform neurons form two separate classes, distinguished by their propensity to fire burst-spikes. We then compare the stimulus-selectivity and rhythmic-synchronization properties among these two different narrow-waveform neurons and the broad-waveform neurons.

## Results

We recorded spikes and Local Field Potentials (LFP) from area V1 in two macaque monkeys (see Methods). Simultaneous recordings were made from 2-10 single electrodes, which had distances to each other between 1 and 3mm. We acutely inserted the electrodes on each recording day and typically positioned them in the superficial layers. During the recordings, we presented drifting-grating stimuli for a duration of 800-1500ms, while the monkeys performed a fixation task. The drifting gratings were centered on the receptive fields (RFs) of the recorded units (≈8° diameter circular aperture). We presented sixteen different stimulus-directions in a random order across trials. Note that statistical parameters are largely described in the figure captions.

### Classification and characterization of neuron types

We used semi-automatic spike-sorting to isolate single units (n=100 in two monkeys). We analyzed the waveforms of the action potentials (APs) to distinguish between narrow- and broad-waveform neurons (Figure 1). For each neuron, we computed the average AP waveform and its peak-to-trough duration (Figure 1A-B). The histogram of peak-to-trough durations was bimodal (Figure 1B). We refer to neurons with long (>0.235ms) and short (<0.235ms) peak-to-trough durations as “broad-waveform” (BW) and “narrow-waveform” (NW) neurons, respectively. The percentage of narrow-waveform neurons was 71% (Figure 1). This replicates the findings of Gur et al. (1999), who observed a similar distribution in all layers of macaque primary visual cortex.

**Figure 1:**
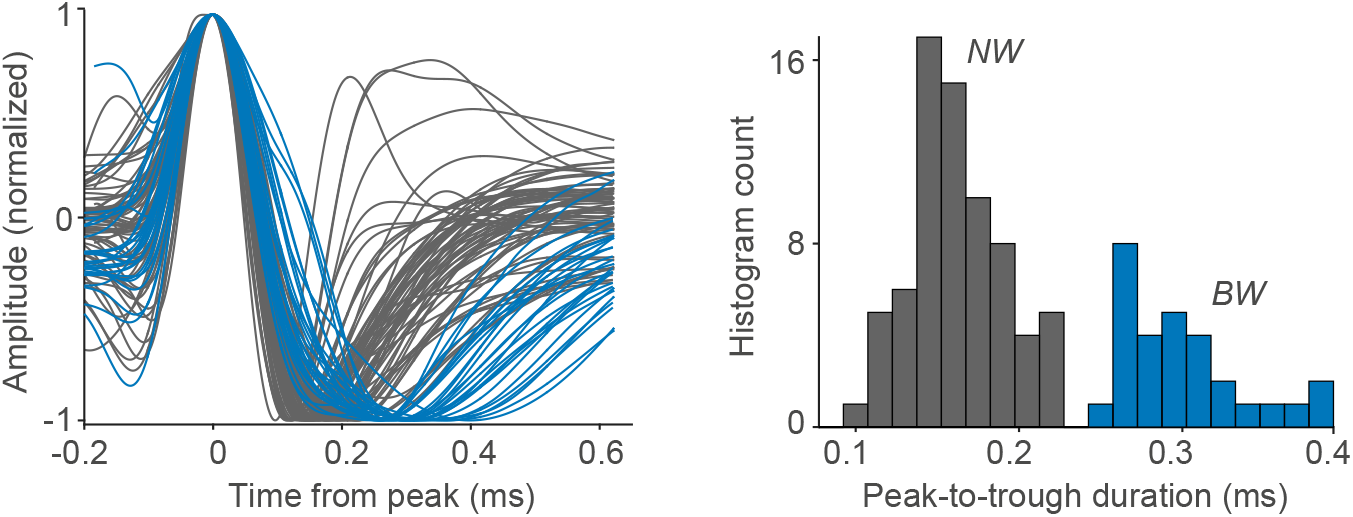
We analyzed the waveforms of action potentials (APs) to distinguish between broad-waveform neurons and narrow-waveform neurons. (A) AP waveforms of the recorded neurons (n=100 cells, two monkeys) as a function of time (ms). We normalized the waveforms between −1 and +1, and alligned them to their respective peaks. Gray and blue traces correspond to narrow-waveform (NW) and broad waveform (BW) APs, respectively. (B) Histogram count of waveform peak-to-trough durations. The distribution of peak-to-trough durations was bimodal (P<0.05; Hartigan’s dip test).

Which cell types do these narrow- and broad-waveform neurons correspond to? Previous studies have distinguished between fast-spiking interneurons and excitatory cells based on the peak-to-trough duration of the AP waveform. This approach is motivated by several considerations: 1) The percentage of inhibitory (GABAergic) interneurons in cortex is small (Hendry et al., 1987; Rudy et al., 2011) 2) Inhibitory interneurons, especially fast-spiking interneurons, typically have narrow-waveform APs (Vinck and Bosman, 2016; Gentet et al., 2012; Senzai et al., 2019; Csicsvari et al., 2003; Miri et al., 2018; Perrenoud et al., 2016; Sirota et al., 2008; McCormick et al., 1985). 3) By contrast, most excitatory neurons have broad-waveform APs (Senzai et al., 2019; Gentet et al., 2012; Vinck et al., 2016; Hasenstaub et al., 2005; Perrenoud et al., 2016; McCormick et al., 1985). 4) In most systems, the percentages of narrow-waveform neurons and inhibitory interneurons are comparable (Senzai et al., 2019; Vinck et al., 2016; Ardid et al., 2015; Miri et al., 2018; Csicsvari et al., 2003; Sirota et al., 2008). Various techniques, including optogenetics, have been used to validate the identification of neuron types based on AP waveform (Senzai et al., 2019; Miri et al., 2018; Vinck et al., 2016; Gentet et al., 2012; Vinck et al., 2016; Bartho et al., 2004). We recorded from single units in L2/3 of mouse V1, and found that the percentage of narrow-waveform neuron is about 9%, consistent with previous studies (Vinck et al., 2015; Niell and Stryker, 2010; Senzai et al., 2019; Vinck et al., 2016). Yet, in case of area V1 in the macaque, the percentage of narrow-waveforms that we observed, ≈70%, was much larger than the expected fraction (20-25%) of GABAergic interneurons (Hendry et al., 1987). This implies that a substantial fraction of NW neurons in macaque V1 might be excitatory rather than inhibitory. Thus, the population of NW neurons should be heterogeneous.

We therefore asked whether this population can be subdivided into distinct neuron types. Neuron types can be identified not only based on the AP waveform, but also based on their characteristic firing patterns. Previous studies have shown that excitatory and inhibitory neurons can be distinguished based on the autocorrelogram of the spike trains (Csicsvari et al., 2003). The autocorrelogram quantifies the likelihood that a neuron spikes at time *t* + *τ*, given that it has spiked at time *t*. Fast-spiking interneurons typically reach a peak in the autocorrelogram at long time-delays >10ms (Vinck et al., 2016; Csicsvari et al., 2003; Senzai et al., 2019; Bartho et al., 2004; Vinck et al., 2013a). This indicates that these neurons have a relatively long refractory-period and do not engage in burst firing. A similar firing behavior can be observed in response to intracellular current injections (Gray and McCormick, 1996; Nowak et al., 2003). By contrast, excitatory neurons often fire in bursts, which results in an early peak in the autocorrelogram (Csicsvari et al., 2003; Vinck et al., 2016; Nowak et al., 2003).

We used a similar approach and computed the autocorrelogram for each of the single neurons. For this purpose, we used the stationary part of the visual stimulus period (>200ms after stimulus onset). We chose this period, because a large number of spikes is needed to reliably estimate the autocorrelogram. We fitted polynomial functions to the autocorrelogram, in order to determine the time at which it reached a global peak (see Methods). In addition, we computed two measures of burst-propensity by comparing the values of the autocorrelogram between short and long time-delays (see Methods). We found that NW neurons formed two separate clusters (Figure 2A-E and Figure S1; see Methods). The first cluster of neurons (“NW–Nonburst”) had a late peak in the autocorrelogram, and did not engage in burst-firing (Figure 2A, C and D-E). These neurons had firing characteristics similar to those of fast-spiking interneuron (see Discussion). The second cluster of neurons (“NW–Burst”) had an early peak in the autocorrelogram, and a high propensity to fire burst-spikes with 2-4ms inter-spike-intervals (Figure 2A-D, Figure 3A). Similar firing patterns in NW neurons were observed in a new-world Capuchin monkey (Figure S3A-B). These characteristics of NW–Burst neurons suggest that they are excitatory, like the BW neurons (Nowak et al., 2003) (we will further address this in the Discussion Section).

**Figure 2:**
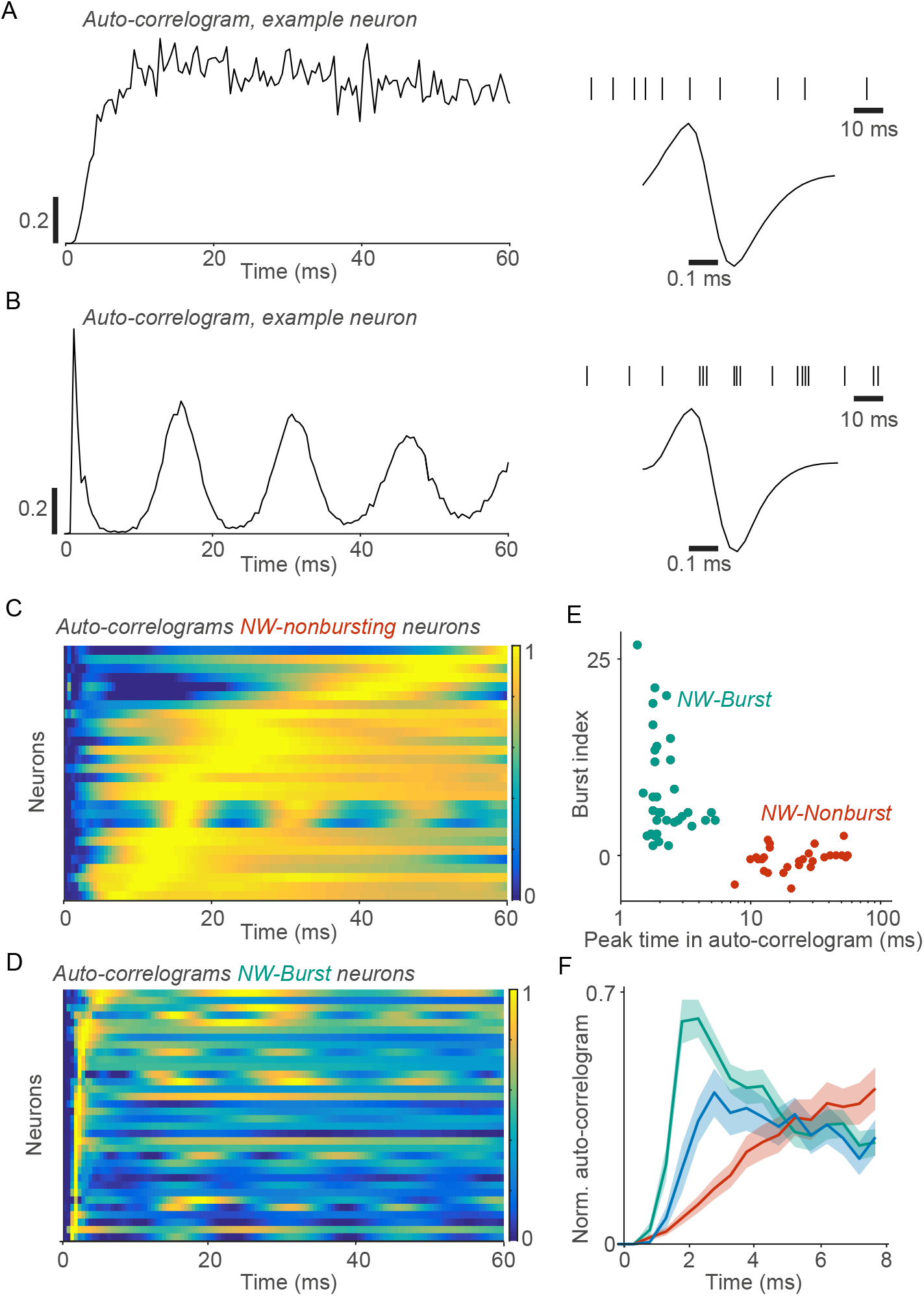
Narrow-waveform neurons formed two non-overlapping clusters, which were distinguished by their bursting-propensity. (A) Example NW neuron. Left: Autocorrelogram as a function of time. The autocorrelogram was normalized between 0 and 1 and computed for the visual stimulation period (>200ms after stimulus onset). Right: Action-potential waveform and an example spike-train during visual stimulation. (B) Similar to (A), but now for another example NW neuron. (C) Autocorrelograms for NW–Nonburst neurons. Each row represents one neuron. Shown are the polynomial fits to the autocorrelograms (see Methods). The rows are sorted according to the time at which the smoothed autocorrelogram reached a global peak (see Methods). (D) Similar to (C), but now for NW–Burst neurons. (E) The time at which the autocorrelogram reached a global peak (within 60ms) vs. burst-propensity (as a Z-score). The burst-propensity measure was constructed by comparing the value of the autocorrelogram between short and long time-delays (see Methods). (F) Mean autocorrelogram for each cell class as a function of time (ms). The autocorrelograms were normalized between 0 and 1 and then averaged across neurons. Shaded regions correspond to standard errors of the mean across neurons. Turqoise, red and blue colors indicate NW–Burst, NW–Nonburst and BW neurons, respectively.

**Figure 3:**
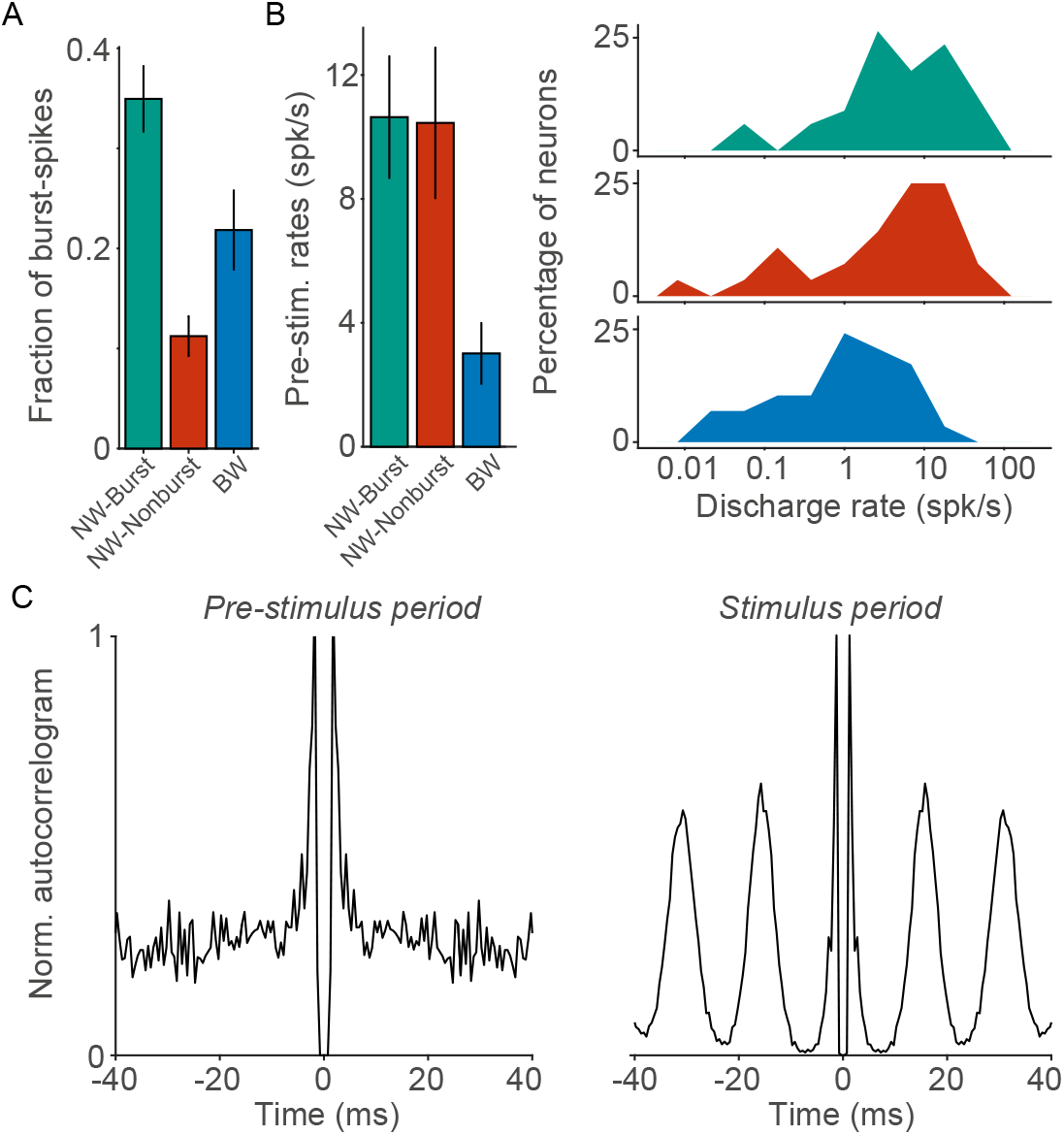
Differences between neuron types in bursting-propensity and in discharge-rates. (A) Mean number of high-frequency burst spikes (subsequent spikes within 6ms intervals) divided by the total number of spikes. This burst-fraction was computed for the visual-stimulation period (> 200ms after stimulus onset). NW–Burst neurons fired more burst-spikes than the other neuron types (BW: P<0.005; NW–Nonburst: P<0.001, permutation test). BW neurons fired more burst-spikes than NW–Nonburst neurons (P<0.05, permutation test). (B) Left: Mean discharge-rates (spikes/sec) during the pre-stimulus period. Right: Distribution of pre-stimulus discharge-rates across neurons. Note that the discharge-rates are shown on a logarithmic scale. NW–Burst and NW–Nonburst neurons had higher rates than BW neurons (NW–Burst vs. BW: *P* < 0.005; NW–Nonburst vs. BW: *P* < 0.001, permutation test). There was no significant difference between NW–Burst and NW–Nonburst neurons (*P* > 0.05, permutation test). (C) Autocorrelograms for an example NW–Burst neuron (same neuron as in Figure 2B). The autocorrelograms were computed both for the pre-stimulus (left) and the stimulus period (right). This neuron had a high burst-propensity in both periods. For a correlation analysis, we refer to Figure S4.

We performed the same analysis for recordings of L2/3 neurons in mouse V1 (see Methods). We used the stationary part of the visual stimulus period during sitting epochs. Thus, the behavioral conditions (drifting gratings, stationary) were comparable between mouse and macaque V1. A subset of the BW neurons showed burst-firing, consistent with (Bartho et al., 2004; Vinck et al., 2016) (Figure S2D-E). Yet, NW neurons showed late peaks in the autocorrelogram, and only one of the NW neurons passed our criterion for NW–Burst neurons, although it had a weak burst-propensity compared to the NW–Burst neurons in macaque V1 (Figure S2D-E). Quantitatively similar results were obtained for L4 in the mouse. Thus, macaque V1 contains a large population of bursting NW neurons that is not found in mouse V1.

We further examined differences in the burst-propensity and discharge rates between NW–Burst, NW–Nonburst and BW neurons in macaque V1. Of the three neuron classes, NW–Burst neurons had the highest burst-propensity (Figure 2F and Figure 3A). A substantial fraction of their spikes had inter-spike-intervals shorter than 6ms (≈35%) and 3.5ms (≈25%) (Figure 3A and Figure S4B). The first spike of the burst had a greater AP amplitude than the subsequent spikes in a burst (*P* < 0.001, T-Test). Conversely, NW–Nonburst neurons rarely fired spikes within 3.5ms intervals (Figure S4B). BW neurons had a higher burst-propensity than NW–Nonburst neurons (Figure 2F and Figure 3A). Yet, the population of BW neurons was heterogeneous (Figure S1) and contained both bursting and non-bursting neurons (Figure S1A). Another defining characteristic of BW neurons was a low discharge-rate (quantified in the pre-stimulus period) as compared to NW–Burst and NW–Nonburst neurons (Figure 3B), similar to mouse V1 (Figure S2C). NW–Burst neurons and NW–Nonburst neurons had similar discharge rates (Figure 3B).

Does the high burst-propensity of NW–Burst neurons depend on visual stimulation, or do they also have a high burst-propensity during spontaneous firing? To investigate this, we computed the autocorrelogram both for the visual stimulation and the pre-stimulus period. The burst-propensity of NW–Burst neurons was significantly correlated between the prestimulus and the stimulus period (Figure 3C and S4D). Their burst-propensity did not significantly differ between the stimulus and pre-stimulus period (Figure S4D). In fact, during the pre-stimulus period, NW–Burst neurons still fired a large fraction of their spikes with inter-spike-intervals shorter than 6ms (Figure S4C). Thus, the high burst-propensity of NW–Burst neurons was not a consequence of visual stimulation, and was also observed during periods of spontaneous firing.

### Orientation selectivity and modulation by grating phase

We proceeded by comparing visual-response properties between the three classes of neurons. We first examined the tuning of discharge rates to the orientation and direction of the drifting-grating stimuli. For each neuron, we computed the orientation-selectivity index (OSI) and the direction-selectivity index (DSI; see Methods). We found that NW–Burst neurons were more orientation-tuned than NW–Nonburst and BW neurons (Figure 4A; see Figure S5 for the two monkeys). By contrast, NW–Nonburst neurons had the weakest orientation- and direction-selectivity of the three neuron types (Figure 4A-B). Thus, the three classes of neurons differed not only in their basic firing properties, but also in their sensory selectivity.

**Figure 4:**
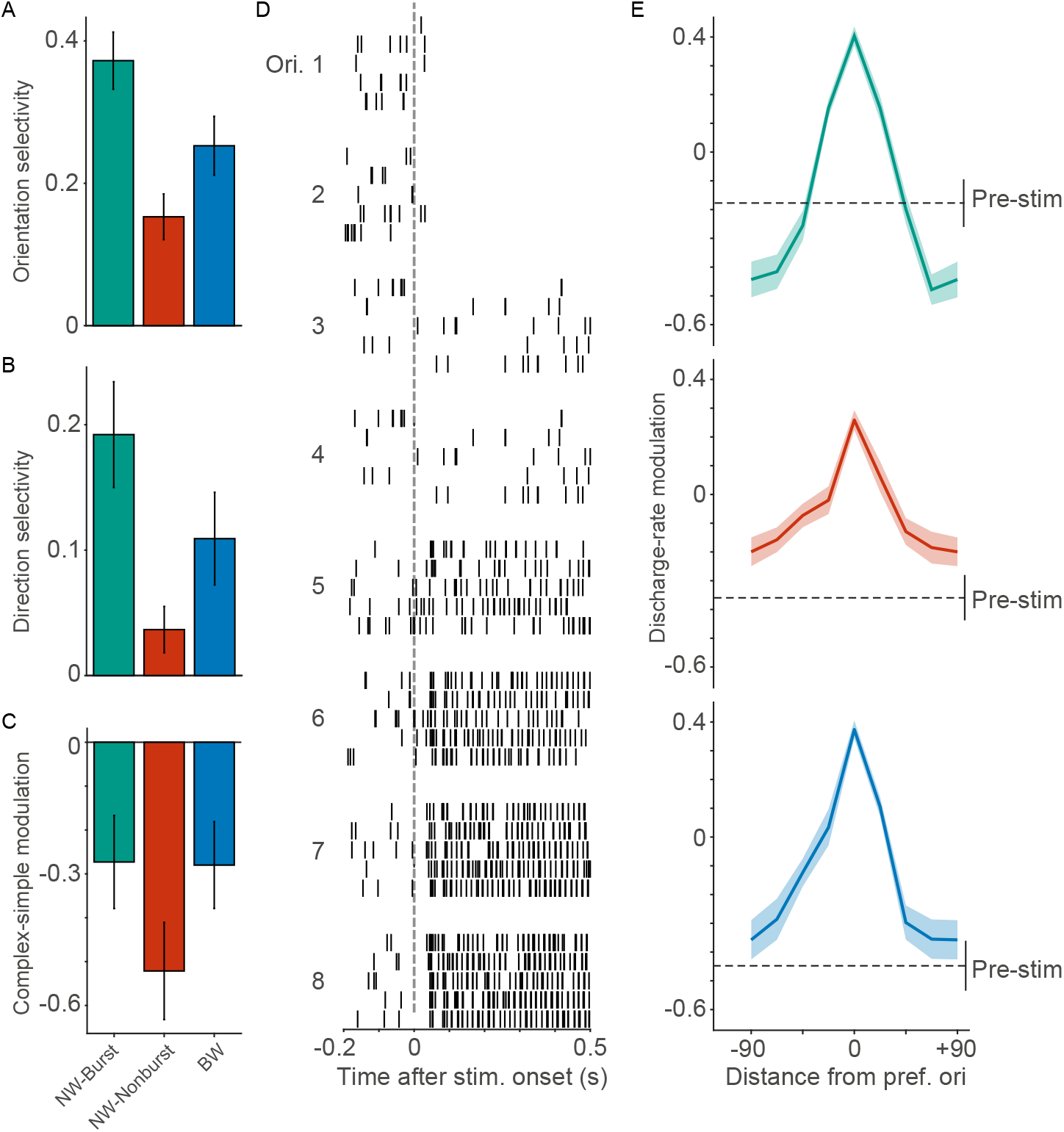
The three neuron types differed in their stimulus-selectivity. NW–Burst neurons were most orientation-selective, and their discharge rates were suppressed for non-preferred stimulus-orientations. (A) Mean orientation-selectivity index (OSI) for the three neuron types. NW–Burst neurons had higher OSI values than NW–Nonburst and BW neurons (NW–Burst vs. NW–Nonburst: P<0.001; NW–Burst vs. BW: *P* < 0.05, permutation test, see Methods). BW neurons had higher OSI values than NW–Nonburst neurons (P<0.05). (B) Mean direction-selectivity index (DSI). NW–Burst and BW neurons had higher DSI values than NW–Nonburst neurons (P<0.001 and P<0.05, respectively; permutation test, see Methods). The DSI values of NW–Burst and BW neurons did not show a significant difference (*P* > 0.05, permutation test). (C) Mean simple vs. complex cell modulation. This modulation was measured as (*F*1 − *F*0)/(*F*1 + *F*0). The F1 represents the modulation of a neuron’s discharge rate by the phase of the drifting-grating (in spikes/sec). The F0 represents the increase in the mean discharge-rate (in spikes/sec). Values higher than zero indicate simple-cell receptive-field properties. BW neurons had higher modulation values than NW–Nonburst neurons (BW vs. NW–Nonburst: *P* < 0.05, permutation test, see Methods). NW–Burst and NW–Nonburst neurons did not significantly differ (P> 0.05, permutation test, see Methods). (D) Example NW–Burst neuron (same neuron as in Figure 2B) We show the spike-raster-plots as a function of time (s) for each of the eight stimulus orientations. The orientations are sorted by the mean discharge-rate. For each orientation, we show the first five stimulus trials. (E) Mean discharge-rate modulation as a function of stimulus orientation. To compute this modulation, we first sorted orientations by the mean change in discharge rate. We then aligned the orientations to the preferred stimulus-orientation. We computed the discharge-rate modulation as (*a* − *b*)/(*a* + *b*). Here, *a* is either the discharge-rate for one of the eight stimulus orientations, or the discharge rate for the pre-stimulus period. The variable *b* represents the mean discharge-rate across orientations. The dashed lines indicate the mean ±1 standard deviation for the pre-stimulus period. For NW–Burst neurons, discharge rates for the pre-stimulus period were higher than for the four worst stimulus orientations (*P* < 0.05, two-sided T-test). For NW–Nonburst neurons, discharge rates were higher for the four worst stimulus orientations than the pre-stimulus period (*P* < 0.05, two-sided T-test). The difference was not significant for BW neurons (*P* > 0.05, T-test). The suppression for NW–Burst neurons was significant as compared to the NW–Nonburst and BW neurons (*P* < 0.001 and *P* < 0.01, permutation test).

Next, we determined the extent to which these neurons had simple or complex receptive-fields. The distinguishing feature of simple V1-cells is that their firing is strongly modulated by the phase of grating-stimuli (Martinez and Alonso, 2003). These cells are predominantly found in L4 (Martinez and Alonso, 2003). By contrast, the discharge rates of complex V1 cells are only weakly modulated by the phase of grating-stimuli (Martinez and Alonso, 2003). These cells are mainly found in L2/3 and L5 (Martinez and Alonso, 2003). For each neuron, we estimated the discharge rate as a function of time by convolving the spike train with a Gaussian kernel (the “spike-density function”; see Methods). This spike-density function was computed only for the preferred stimulus-orientation. We then fitted a sinusoid to this spike-density function and computed a modulation measure as *M* = (*F*1 − *F*0)/(*F*0 + *F*1) (see Methods). Here, *F*1 is the amplitude of the sinusoid (corrected for estimation-bias) and *F*0 is the mean elevation of firing above baseline (see Methods). Values of *M* below and above zero indicate complex-cell and simple-cell modulation, respectively. In line with previous work, we observed a bimodal distribution of modulation values (Hartigan’s dip test: *P* < 0.05). Most neurons had complex receptive-field properties, which is consistent with the placement of our electrodes in superficial layers (Figure 4C; *M* < 0 for 76% of NW–Burst neurons; NW–Nonburst: 85%, BW: 70%). The discharge rates of BW neurons were more strongly modulated by the phase of the drifting gratings than the rates of NW–Nonburst neurons (Figure 4C). NW–Burst neurons mainly had complex RF properties, and did not significantly differ from NW–Nonburst and BW neurons (Figure 4C). In sum, neurons mainly had complex RFs, and the RFs of NW–Nonburst neurons tended to be more like the complex type.

We found that NW–Burst neurons were more selective for stimulus orientation than the other cell classes. We had expected the opposite finding, because of their high spontaneous discharge-rates. One explanation could be that these neurons receive broadly-tuned and strong inhibitory inputs restricted to the period of visual stimulation (Shapley et al., 2003; Isaacson and Scanziani, 2011). This would decrease their excitability, and thereby sharpen their orientation tuning. To examine this, we compared the neurons’ discharge rates between the stimulus and the pre-stimulus period. We observed that some of the NW–Burst neurons were highly suppressed at their non-preferred stimulus-orientation (Figure 4D). To study this phenomenon at the population level, we determined the mean discharge rates for each stimulus orientation separately. We found that the discharge rates of NW–Burst neurons were suppressed below baseline levels for non-preferred orientations (Figure 4D-E). This pattern was not observed for NW–Nonburst and BW neurons (Figure 4E). Thus, NW–Burst neurons encoded stimulus orientation not only with an increase, but also with a decrease in firing relative to the pre-stimulus period. This finding is consistent with the study of Shapley et al. (2003), who found that some V1 neurons show suppression at non-preferred stimulus orientations. Together, this indicates that NW–Burst neurons receive a strong inhibitory drive during the visual-stimulation period, which can sharpen the tuning of these neurons (Shapley et al., 2003).

### Differences in rhythmic firing

The way that V1 neurons transmit stimulus information via horizontal and feedforward connections likely depends on the synchronization of their responses (Fries, 2015; König et al., 1996; Salinas and Sejnowski, 2000). Many visual stimuli induce prominent 30-80Hz gamma-oscillations in primate V1 (Fries, 2009; Vinck and Bosman, 2016; Lima et al., 2010). However, the contributions of different neuron types to gamma-synchronization in primate V1 remain largely unknown.

To investigate this, we compared the strength of spike-LFP phase-locking between the three neuron types. For each spike, we determined its phase relative to the LFP recorded from the other electrodes (see Methods). After computing the spike-LFP phases, we estimated phase-locking using the pairwise-phase-consistency (PPC) measure (Vinck et al., 2012). The PPC is not affected by mean discharge rates and history effects like bursting (see Methods) (Vinck et al., 2012). Thus, it can be used to compare neuron types with different levels of discharge rates and burst-propensity. Gamma-phase locking is illustrated for an example NW–Burst neuron in Figure 5. We found that all three neuron types were phase-locked to LFP gamma oscillations within a narrow frequency-range (Figure 6A-C and Figure S6A). Yet, NW–Burst neurons showed much stronger gamma-phase locking than BW and NW–Nonburst neurons (≈2.5-fold difference; p<0.05, permutation test). This pattern was consistent across the two monkeys (Figure 6A and Figure S6A-B). This is consistent with the presence of rhythmic sidelobes in the autocorrelogram of NW–Burst neurons (Figure 2D). Note that a small subset of NW–Nonburst neurons exhibited rhythmic sidelobes in the autocorrelogram (Figure 2C). In sum, we find that the class of NW–Burst neurons are strongly gamma-synchronizated as compared to the other cell classes.

**Figure 5:**
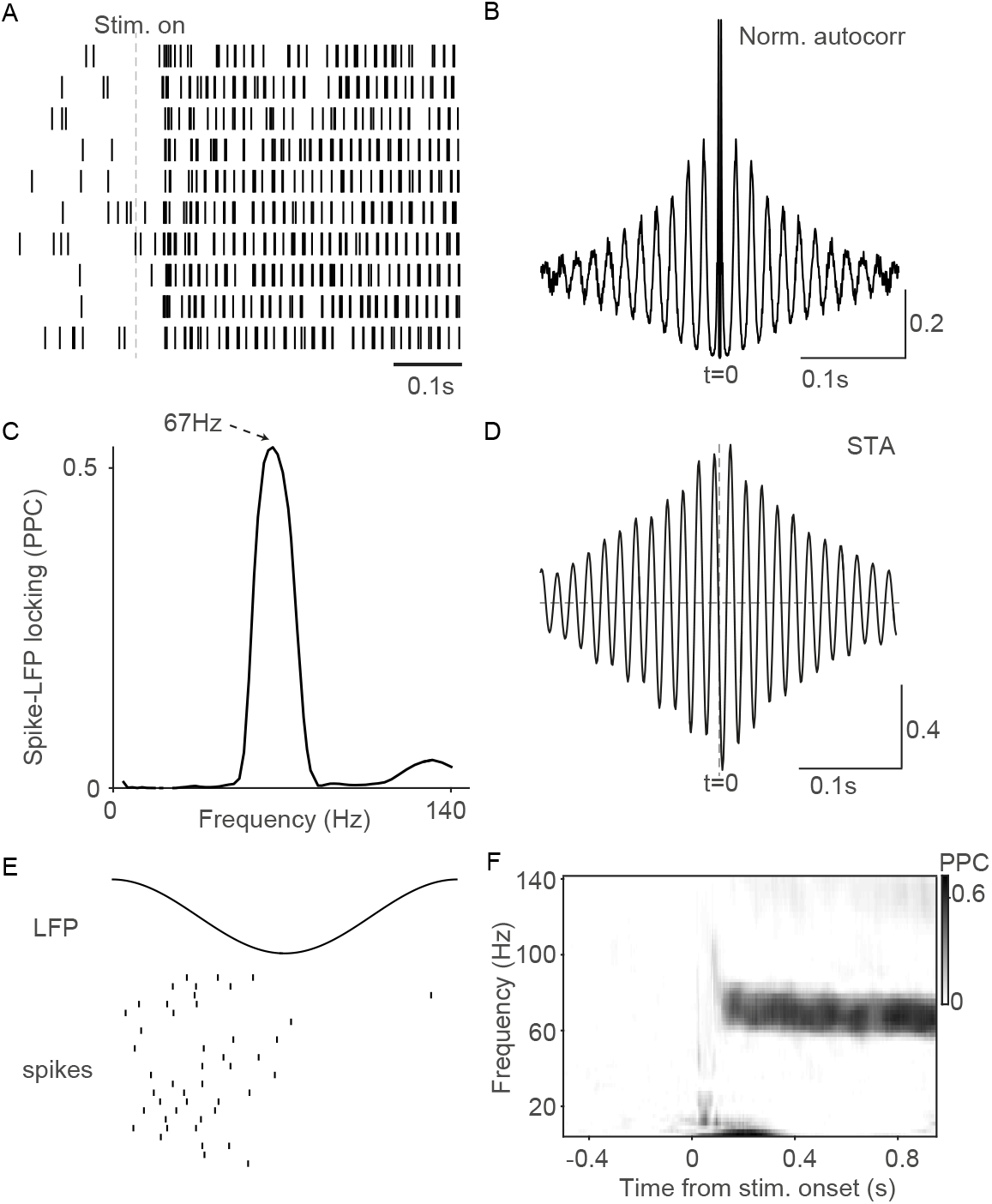
Spike-LFP gamma phase-locking for an example NW–Burst neuron (same neuron as in Figure 2B). (A) Raster-plot of spikes recorded in response to a drifting grating with the neuron’s preferred stimulus-direction. Shown are the first 10 trials. (B) Normalized (between 0 and 1) autocorrelogram during the visual stimulation period (> 200ms after stimulus onset). The autocorrelogram had oscillatory side-lobes at a period of the gamma cycle (≈ 15ms), which indicates gamma-rhythmic firing. (C) Spectrum of spike-LFP gamma-phase locking as a function of frequency (Hz). Spike-LFP phase-locking was estimated with the pairwise-phase-consistency (PPC; see Methods). (D) Spike-triggered average (STA) of the LFP as a function of time (s). Before computing the STA, we Z-scored the LFP signals. The STA has oscillatory side-lobes at the period of a gamma cycle, similar to the autocorrelogram. (E) Raster-plot of spikes as a function of gamma phase. Each row represents a different gamma-cycle. We show a selection of gamma cycles that had a duration of 15ms and that were detected during the trials in which the preferred stimulus-direction was presented. Spikes were clustered at the falling phase of the LFP gamma cycle. (F) Spike-LFP gamma phase-locking (PPC) as a function of frequency (Hz) and time (s). The neuron showed gamma-phase locking shortly after the onset of the visual stimulus. This gamma-locking was sustained throughout the stimulus period.

**Figure 6:**
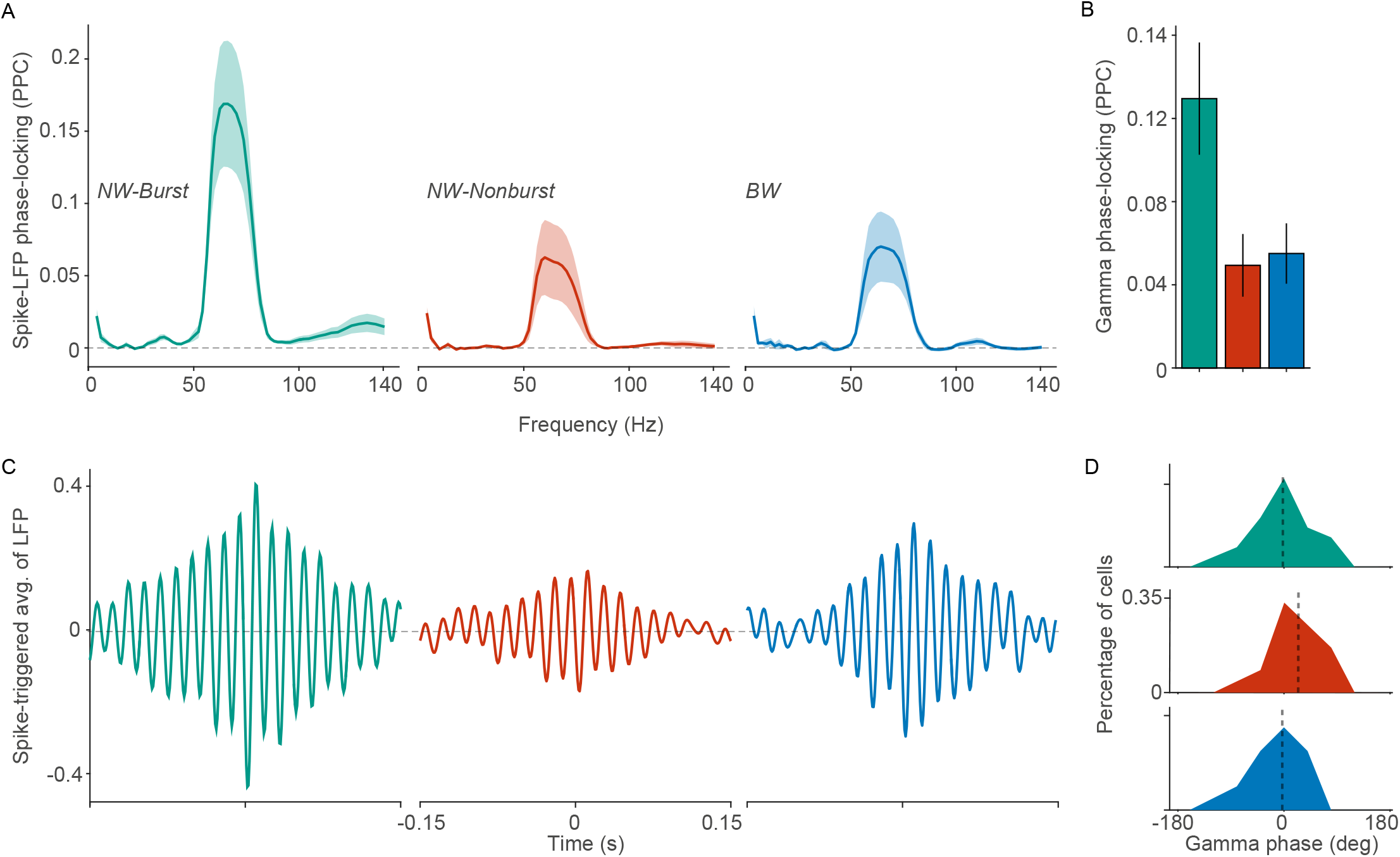
NW–Burst neurons were more gamma phase-locked than NW–Nonburst and BW neurons. (A) Mean spike-LFP phase-locking (PPC) values as a function of frequency (Hz) for monkey J. We computed the spike-LFP phases using a short-term Fourier-transform with 9 cycles per frequency and a Hann taper (see Methods). For monkey L, see Figure S6A. (B) Mean gamma phase-locking (PPC) across all recorded neurons from both macaques. In this case, we computed the spike-LFP phases by determining the peaks and troughs of the band-pass filtered LFP signals (see Methods). NW–Burst neurons had higher gamma phase-locking than NW-Nonbursting and BW neurons (2.62 and 2.35 times stronger; both comparisons *P* < 0.01, permutation test; see Methods). BW and NW–Nonburst neurons were not significantly different (*P* > 0.05). (C) Bottom: Mean spike-triggered-average (STA) of the LFP as a function of time around the spike. We computed the mean STAs by averaging across neurons. Before computing the STA, we Z-scored the LFP signals. Note that the STA is biased as a function of the spike count, such that lower spike counts yield STAs with a smaller amplitude. The STAs exhibit oscillatory side-lobes, whose amplitude decays slowly over time. This is consistent with the narrow-band character of spike-LFP phase-locking. (D) Distribution of preferred gamma-phases across neurons. Shown is the difference between the preferred gamma-phases of the single units and the same-site MUAs (see Methods). The spikes of NW–Bursting neurons were significantly delayed relative to the same-site MUA spikes (95% confidence intervals: 11.7-84.6 degrees; difference to BW and NW–Burst neurons: *P* < 0.05, permutation test).

Gamma oscillations are often studied in the later part of the stimulus period, which avoids the non-stationary, transient response that occurs shortly after stimulus onset. The way in which we computed the spike-LFP phases, allowed us to study gamma phase-locking in an earlier part of the stimulus period (100-250ms after stim. onset; see Methods). Also in this early phase of the stimulus-period, NW–Burst neurons were more gamma phase-locked than NW–Nonburst and BW neurons (Figure S6C). We observed similar levels of phase-locking in the early and late phase of the stimulus period (>200ms after stimulus onset). Hence, gamma-phase locking may play a functional role in “early vision”.

Many factors can influence gamma-phase locking, for example the laminar position of the electrode and the state of the animal (van Kerkoerle et al., 2014; McGinley et al., 2015b; Herculano-Houzel et al., 1999; Livingstone, 1996; Vinck et al., 2015; Senzai et al., 2019; McGinley et al., 2015a; Zagha et al., 2013; Xing et al., 2012; Herculano-Houzel et al., 1999; Gray et al., 1992; Besserve et al., 2015). These factors may have contributed to the observed differences between neuron types. To investigate this, we used a similar method as in Vinck et al. (2013a): For each single unit, we constructed a “same-site MUA” signal. This signal contained all the remaining spikes that were recorded from an electrode, after excluding the activity of the isolated single units. For each same-site MUA, we computed the phase-locking of MUA spikes to LFP oscillations (PPC). This allowed us to make a comparison between same-site-MUAs corresponding to NW–Burst, NW–Nonburst and BW neurons. We did not observe a significant difference in phase-locking between the same-site-MUAs of NW–Burst, NW–Nonburst and BW neurons (all comparisons *P* > 0.05, permutation test). Moreover, NW–Burst neurons had much higher gamma phase-locking than the same-site-MUA (Figure S6B). This was not the case for NW–Nonburst and BW neurons (Figure S6B). These results suggest that other factors like electrode position and behavioral state did not explain the differences in gamma-locking between neuron types.

Discharge rates and burst-propensity could also have influenced gamma phase-locking. Note that the PPC measure removes biases due to spike count and/or history effects like bursting (Vinck et al., 2012). Nevertheless, we wanted to examine whether burst-propensity and discharge rates correlate with phase locking. For each neuron type, we correlated the prestimulus discharge rates with the gamma phase-locking during the visual stimulation period. We found no significant correlation for any of the three neuron types (NW–Burst: *ρ*=−0.2976, NW–Nonburst: −0.1174, BW: 0.0220, *P* > 0.05 for all neuron types). Likewise, burst-propensity (measured during the stimulus period) was not significantly correlated with gamma phase-locking (NW–Burst: *ρ* = −0.1202, BW: −0.0098, *P* > 0.05; note that NW–Nonburst neurons lacked the necessary bursts for this analysis). In fact, the high burst-propensity of NW–Burst neurons may have decreased their phase-locking: Suppose that a neuron fires in bursts, and that the first spike of a burst is always precisely timed to the onset of the gamma cycle. The other spikes of the burst would then occur at later gamma phases, thereby reducing phase-locking. To investigate this possibility for NW–Burst neurons, we detected all bursts and computed the gamma phase-locking for the set of spikes that came first in a burst. Indeed, we confirmed that the first spikes of the burst were (≈ 2-fold) more gamma phase-locked than all spikes together (Figure S6E). This means that the gamma phase-locking of NW–Burst neurons was even more precise for the first spikes of their bursts.

Models of gamma oscillations make specific predictions about the relative timing of different neurons types within the gamma cycle (Buzsáki and Wang, 2012; Wang, 2010; Tiesinga and Sejnowski, 2009; Börgers and Kopell, 2005, 2008) We therefore asked whether the three different neuron-types fired at similar phases of the gamma cycle, or whether they fired in a specific sequence. Previous studies on gamma have found that fast-spiking interneurons fire with a short delay after excitatory neurons, which is consistent with PING models of gamma generation (Vinck et al., 2013a; Csicsvari et al., 2003; Hasenstaub et al., 2005). In PING models, a rise in excitatory-cell firing triggers an increase in inhibitory-cell firing at a short delay, leading to a subsequent decrease in the firing of excitatory cells (Buzsáki and Wang, 2012; Tiesinga and Sejnowski, 2009; Börgers and Kopell, 2005, 2008; Wang, 2010). Whether these dynamics can be observed in area V1 remains unknown. For each neuron, we determined the gamma phase at which its discharge rate reached a peak (the “preferred gamma-phase”; see Methods). A potential caveat in the analysis of spike-LFP phases is that there is substantial variability in the phase of firing across recording sites (Vinck et al., 2013a; Livingstone, 1996; van Kerkoerle et al., 2014). This is caused by many factors, especially the stimulus drive and the laminar position of the unit or LFP (Livingstone, 1996; van Kerkoerle et al., 2014; Vinck et al., 2013a, 2010a). To remove this variability, we computed the preferred gamma-phase for each of the same-site-MUAs (see Methods). We then computed the difference between the preferred gamma-phases of the single unit and of its corresponding same-site-MUA. The distributions of phase differences is shown in Figure 6D. We found that the firing of NW–Nonburst neurons was delayed relative to the same-site MUA (Figure 6D). By contrast, the phase of NW–Burst and BW neurons did not differ relative to the same-site MUA (Figure 6D). NW–Nonburst neurons fired significantly later in the gamma cycle than NW–Burst and BW neurons (*P* < 0.05, permutation test; see Methods). This is consistent with the idea that NW–Nonburst neurons correspond to fast-spiking inhibitory interneurons, and fire with a short delay after excitatory neurons.

### Dependence of the discharge rate on the gamma phase

In the analysis of orientation tuning presented above, we observed that the firing of NW–Burst neurons was suppressed for the non-preferred stimulus orientation (Figure 4D-E). This suggests that these neurons receive a very strong inhibitory drive during the visual stimulation period. We wondered whether a similar suppression occurs during the non-preferred LFP gamma phases of NW–Burst neurons. The non-preferred gamma phase of a neuron is defined as the gamma phase at which its discharge rate reaches a minimum. We determined neuronal discharge-rates as a function of the gamma phase as follows: We first binned the LFP gamma cycle in 40 nonoverlapping phase bins. We then determined the amount of time that the LFP spent in a given gamma phase-bin. This was equal to 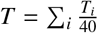, where *T_i_* is the duration of the *i*-th recorded gamma cycle. The discharge rate in the *k*-th phase bin then equaled *N_k_*/T, with *N_k_* the number of recorded spikes in that phase bin. The discharge rate as a function of gamma phase for an example NW–Burst neuron is shown in Figure 7A-B. This neuron was virtually silent at its non-preferred gamma phase (Figure 7B). We found that on average, the discharge rates of NW–Burst neurons were suppressed during non-preferred phases of the gamma cycle (Figure 7C). This suppression was not found in NW–Nonburst and BW neurons (Figure 7C).

**Figure 7:**
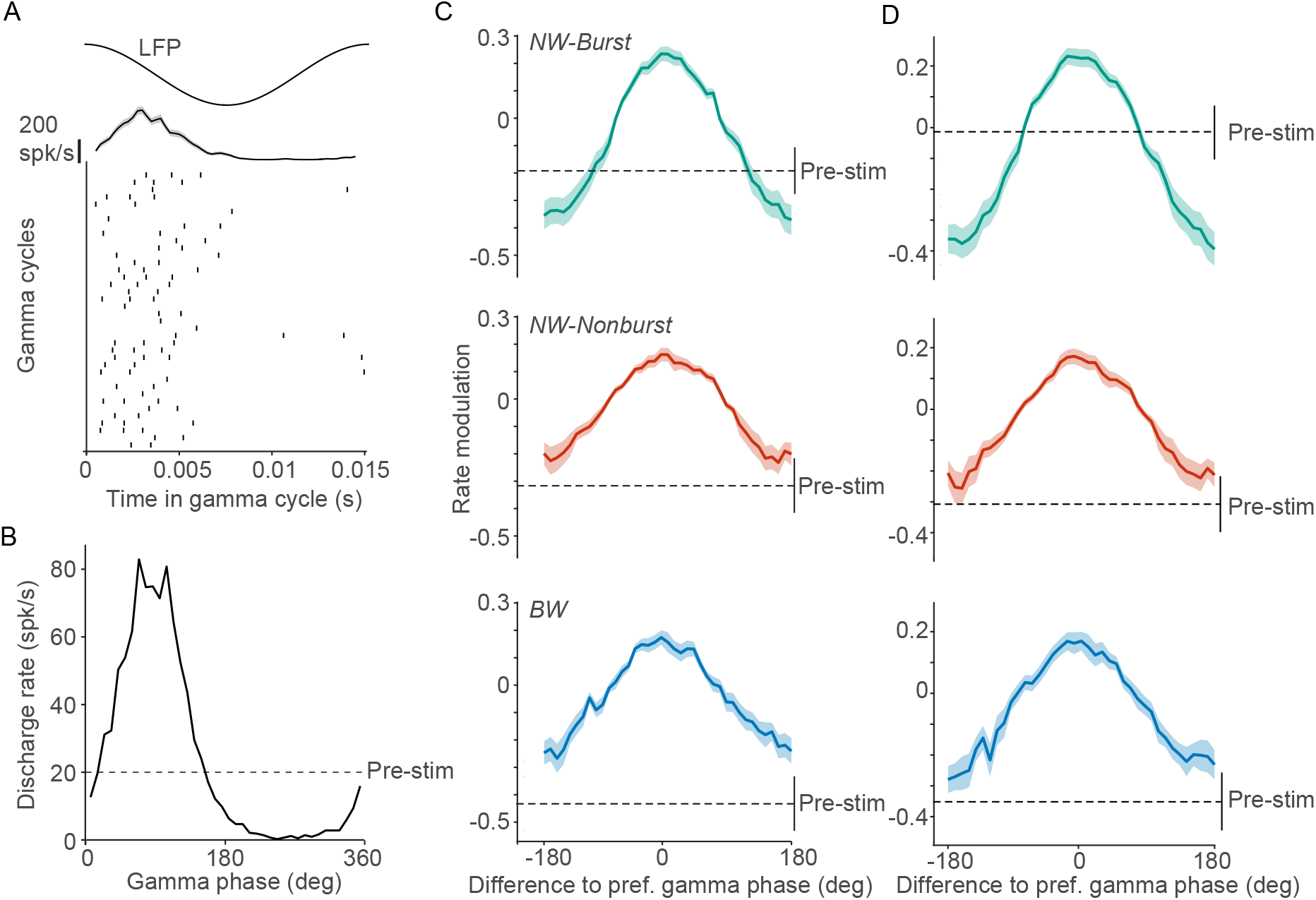
Discharge rates of NW–Burst neurons were suppressed at their non-preferred gamma phase. (A) Example NW–Burst neuron (same neuron as in Figure 2B). Raster-plot of spikes in the gamma cycle. Each row represents a different gamma cycle, aligned to the LFP peak. We show a selection of gamma cycles that had a duration of 15ms and were detecte during trials in which the preferred stimulus-direction was presented. At the top, we show the instantaneous discharge-rate as a function of the gamma phase. (B) Mean discharge rate (spikes/sec) as a function of the gamma phase (degrees) for the example neuron shown in (A). We computed the mean discharge rate by averaging across all stimulus trials. The dashed line represents the neuron’s pre-stimulus discharge-rate. (C) Mean modulation of the discharge rate as a function of the gamma phase (degrees). For each neuron we computed the preferred gamma-phase. The x-axis represents the phase relative to the preferred gamma-phase (degrees). The modulation of the discharge rate was computed as (*a* − *b*)/(*a* + *b*). Here, *a* was the discharge-rate for either one of the 40 gamma phases, or for the pre-stimulus period. The variable *b* represents the mean discharge-rate across gamma phases. The dashed lines indicate the mean for the pre-stimulus period. The vertical line on the right indicates ±1 standard error of the mean for the pre-stimulus period. NW–Burst neurons were significantly suppressed relative to baseline (*P* < 0.05, T-Test). The difference between NW–Burst and NW–Nonburst and BW neurons was significant (*P* < 0.05, permutation test; see Methods). (D) Similar to (C), but now computing the discharge rates in a different way. For each *k*-th gamma phase bin, we determined whether the last spike had occurred at least 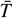 milliseconds before the gamma phase bin, where 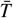 was the average period of the gamma cycle. We then counted the number of phase bins for which this was the case, and computed the total amount of time spent in that gamma phase bin, *T*. The discharge rate then equaled *N_k_*/*T*, where *N_k_* was the number of spikes in those gamma phase bins (i.e. occurring after a pause of no firing for 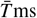). NW–Burst neurons were significantly suppressed relative to baseline (*P* < 0.001, T-Test). The difference between NW–Burst and NW–Nonburst and BW neurons was significant (*P* < 0.001).

One possible explanation for the suppression of discharge rates during the non-preferred gamma phase is a strong inhibitory input drive arriving at that phase. An alternative explanation could be that NW–Burst neurons are silent at their non-preferred gamma phases, because they have already fired at an earlier phase. This could suppress firing at later gamma phases, if these phases occur during the neuronal refractory period. To distinguish between these two scenarios, we computed discharge rates in an alternative way: For each *k*-th gamma phase bin, we determined whether the last spike had occurred at least 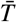 milliseconds before the gamma phase bin, where 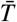 was the average period of the gamma cycle. We then counted the number of phase bins for which this was the case, and computed the total amount of time spent in that gamma phase bin, *T*. The discharge rate then equaled *N_k_*/*T*, where *N_k_* was the number of spikes in those gamma phase bins (i.e. occurring after a pause of no firing for 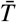ms). We found that the suppression at the non-preferred gamma phase for NW–Burst neurons was even more pronounced when the discharge rates were calculated in this way (Figure 7D). Hence, firing suppression at the non-preferred gamma phase was not due to the neuronal refractory period.

In sum, a unique feature of NW–Burst neurons is that their firing is decreased below pre-stimulus levels during the nonpreferred gamma phase. This is a surprising finding, considering that visual stimuli should increase the excitatory drive to NW–Burst neurons. It thus suggests that these neurons receive a strong inhibitory input drive at the non-preferred gamma phase. Notably, in Figure 4D-E we showed a similar suppression of NW–Burst neurons for non-preferred stimulus-orientations.

### Relationship between gamma and orientation tuning

We observed differences in both orientation selectivity and gamma-synchronization between neuron types. In particular, we found that NW–Burst neurons have both high orientation-selectivity and gamma phase-locking as compared to other neuron types. Previous work has suggested a close relationship between these two aspects of V1 activity (Friedman-Hill et al., 2000; Vinck et al., 2010a; Womelsdorf et al., 2012; Maldonado et al., 2000). We previously reported a positive correlation between gamma phase-locking and orientation-selectivity across neurons (Womelsdorf et al., 2012). It is likely that this positive correlation was partly due to the strong phase-locking and orientation-selectivity of NW–Burst neurons. We therefore revisited this analysis, and examined the correlations for the three neuron classes separately. NW–Burst neurons showed a highly positive correlation between orientation selectivity (OSI) and gamma phase-locking values (Figure 8A-B; *P* < 0.001, bootstrap test). This correlation was weaker and not significant in the other two neuron types (Figure 8A-B; *P* > 0.05, bootstrap test; NW–Burst vs. BW and NW–Nonburst, *P* < 0.05, permutation test). NW–Burst neurons also showed a negative correlation between gamma phase-locking and the extent to which their firing was modulated by the phase of the grating (i.e. simple vs. complex) (R=-0.27, *P* < 0.05, Bootstrap-test). Thus, NW–Burst neurons with more complex receptive fields and stronger orientation tuning were also more gamma phase-locked.

**Figure 8:**
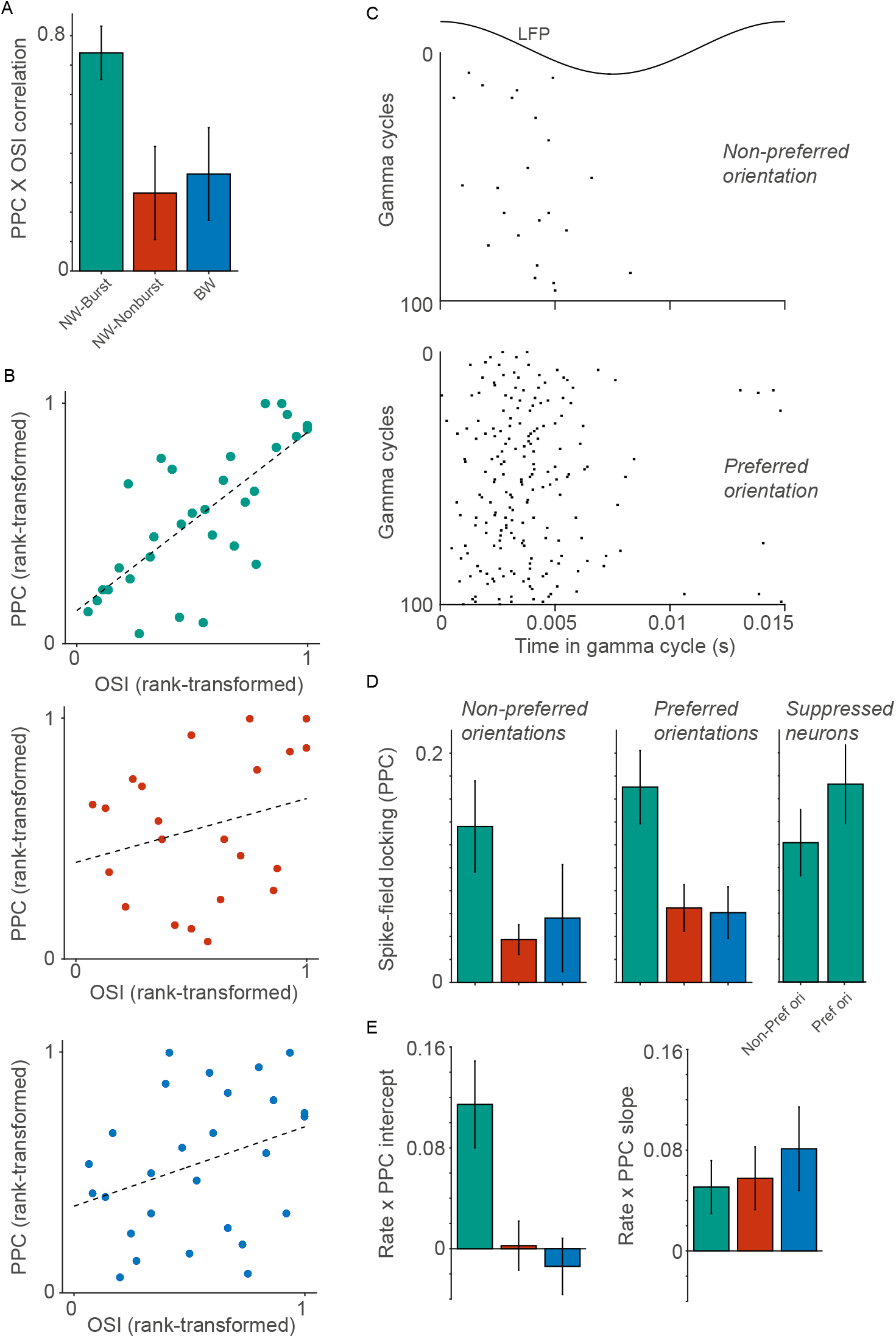
The gamma phase-locking of NW–Burst neurons was strongly correlated with their orientation-selectivity. NW–Burst neurons were also gamma phase-locked for non-preferred stimulus orientations. (A) Rank-correlation between orientation-selectivity index (OSI) and spike-LFP phase-locking (PPC). We first rank-transformed the OSI and PPC values and normalized the rank scores between 0 and 1. We then pooled the normalized ranks across the two monkeys, and computed the Pearson’s correlation coefficient. Because of the rank-transformation, this coefficient is comparable to the Spearman’s rho correlation. We computed the standard errors of the mean by boostrapping over neurons. Correlations were significantly higher for NW–Burst than NW–Nonburst and BW neurons (*P* < 0.05, permutation test). (B) Correlation between OSI and PPC for NW–Burst neurons (top), NW–Nonburst neurons (middle), and BW neurons (bottom). We show the normalized ranks of the OSI (x-axis) and the PPC (y-axis). (C) Raster-plot of spikes as a function of gamma phase for an example NW–Burst neuron (same neuron as in Figure 2B). Each row represents a different gamma cycle, aligned to the LFP peak. We show a selection of gamma cycles that had a duration of 15ms. Top and bottom: Gamma cycles that were detected during trials in which a non-preferred (top; 4th orientation from the top in Figure 4D) or preferred (bottom) stimulus-orientation was presented. (D) Left: Spike-LFP phase-locking (PPC) for the three stimulus orientation at which the neurons’ discharge rates were the lowest. For (D), we only included neurons that were significantly orientation-tuned. Middle: PPC for the stimulus orientation at which the neurons’ discharge rates were the highest. Right: Same analysis, but now for NW–Burst neurons whose firing rates were suppressed relative to baseline for the three non-preferred stimulus orientations. There were no significant differences between preferred and non-preferred stimulus orientations for all comparisons (*P* > 0.05). (E) Left: Mean intercept for the regression of spike-LFP gamma phase-locking (PPC) onto discharge rate. For each neuron, we computed the discharge rate and PPC for each stimulus direction separately. We normalized the discharge rates by dividing by the maximum discharge rate. We then predicted the PPC values from the discharge rates. The intercept estimates the PPC as the normalized discharge rate converges to 0. Right: Mean slope for the regression of PPC onto discharge rate. The slope represents the change in PPC that would occur as the normalized rate increases from 0 to 1.

Previous experimental and computational work has suggested that gamma oscillations depend on stimulus drive (Ray and Maunsell, 2010; Peter et al., 2019; Bartos et al., 2007; Gray et al., 1990; Gray and Di Prisco, 1997; Roberts et al., 2013; Jia et al., 2011, 2013; Chalk et al., 2010). For example, gamma-amplitude increases with luminance-contrast (Henrie and Shapley, 2005; Ray and Maunsell, 2010; Gray and Di Prisco, 1997; Roberts et al., 2013), and it can be abolished by stimulus adaptation (Peter et al., 2019). What explains this drive-dependence? It could be that sufficient drive to the entire neuronal population is required for the emergence of gamma-synchronization. An alternative possibility is that a single neuron can only entrain to gamma oscillations when its membrane potential is sufficiently depolarized. Because luminance-contrast modulates the firing of most neurons in V1, its manipulation cannot be used to distinguish between these two scenarios. However, when we present a grating with a specific stimulus-orientation, some neurons will be strongly driven, and other neurons in the same hypercolumn will be suppressed. For example, the NW–Burst neuron shown in Figure 8C was strongly suppressed during the presentation of its non-preferred stimulus orientations (Figure 4B). Yet, it still showed gamma phase-locking for non-preferred orientations (Figure 8C), suggesting that a given neuron can be strongly gamma-locked even if it is not strongly driven.

To investigate this at the population level, we used two approaches. In the first approach, we compared the gamma phase-locking between a neuron’s best (i.e. preferred) stimulus orientation and its three worst (i.e. non-preferred) orientations (Figure 8D). The best orientation was defined as the orientation at which its discharge rate was maximum. Note that we pooled the three worst orientations to obtain a sufficient amount of spikes to reliably estimate spike-LFP locking values. Surprisingly, we found that for all three neuron types, gamma phase-locking was not higher for the preferred than for the non-preferred stimulus orientations (T-Test: *P* > 0.05, Figure 8D). However, it could be that there was some stimulus drive for the nonpreferred stimulus orientations, which could have resulted in gamma phase-locking. We therefore selected the NW–Burst neurons whose discharge rates were suppressed below baseline for their three worst stimulus-orientations. Also for these neurons, we found that gamma phase-locking did not significantly differ between non-preferred and preferred stimulus orientations (T-Test: *P* > 0.05, Figure 8D). Thus, gamma phase-locking was largely independent of stimulus orientation, and remained strong even when discharge rates were suppressed below baseline levels. This suggests that a neuron’s gamma phase-locking depends on sufficient drive to the entire network, but not on the excitatory drive to that individual neuron.

However, this analysis may have been insensitive, because we ignored a subset of orientations. In the second approach, we predicted the PPC values from the discharge rate across all the 16 stimulus directions (Figure 8E). For each neuron, we computed the discharge rates for the 16 different stimuli separately. The firing rates were normalized by dividing by the maximum discharge rate. Thus, normalized discharge rates were bounded by 0 (a discharge rate of zero) and 1 (the maximum discharge rate). We then predicted the 16 gamma phase-locking values from the 16 normalized discharge-rates (see Methods). The slope of the regression models indicated the change in PPC as the normalized discharge rate increased from 0 to 1. The intercept indicated the level of phase-locking that would be expected if the discharge rates converged to zero. We found that gamma phase-locking was positively related to discharge rates for all three neuron types (*P* < 0.05, T-Test; Figure 8E). However, the regression intercepts were significantly higher than zero for NW–Burst neurons, but not for BW and NW–Nonburst neurons (T-Test: *P* < 0.001; Figure 8D). This indicates that NW–Burst neurons exhibited strong gamma phase-locking even when their discharge rates approached zero (Figure 8E). This result is consistent with the direct comparison of preferred and non-preferred orientations shown in Figure 8D. In addition, this analysis shows that gamma phase-locking increased as a function of stimulus-driven discharge rates.

## Discussion

The contributions of distinct neuron types to synchronization and stimulus selectivity in primate V1 remain largely undetermined. We recorded spikes and LFPs from awake macaque V1 and analyzed AP waveforms to distinguish between narrow- and broad-waveform neurons. Narrow- and broad-waveform APs are commonly used to identify fast-spiking interneurons and excitatory cells, respectively. Surprisingly, most neurons in macaque V1 had a *narrow* waveform (NW) AP. This suggests that the population of NW neurons is heterogeneous and contains excitatory neurons. By analyzing firing statistics, we showed that NW neurons form two separate clusters: NW–Burst neurons had early peaks in the autocorrelogram and a high burst-propensity. Conversely, NW–Nonburst neurons had late peaks in the autocorrelogram and a low burst-propensity. As we will discuss further below, these and other properties suggest that NW–Burst and NW–Nonburst neurons correspond to (bursty) excitatory neurons and fast-spiking interneurons, respectively. Comparisons between NW–Burst, NW–Nonburst and BW neurons revealed major differences in firing properties, visual selectivity and oscillatory synchronization. Most strikingly, we found that NW–Burst neurons were most strongly phase-locked to 30-80Hz gamma oscillations and were most stimulus-selective. Furthermore, we found signatures of strong rhythmic inhibition in NW–Burst neurons, suggesting that they interact with GABAergic interneurons to act as excitatory pacemakers for the V1 gamma rhythm.

### Classification of neuron types

We distinguished three neuron types based on AP waveform and firing properties. One class of neurons had broad waveforms (BW). Previous studies identified these as excitatory (Rudy et al., 2011; Gentet et al., 2012; Nowak et al., 2003; Vinck et al., 2016; Senzai et al., 2019; Miri et al., 2018; Perrenoud et al., 2016). This inference has been validated based on optogenetics, electrophysiology and cell-targeted recordings (Nowak et al., 2003; Senzai et al., 2019; Gentet et al., 2012; Vinck et al., 2016; Miri et al., 2018; Hasenstaub et al., 2005; Perrenoud et al., 2016). Consistent with previous studies, we found that BW neurons had low spontaneous discharge-rates, and that a subset of BW neurons exhibited burst-firing (Vinck et al., 2016; Senzai et al., 2019; Bartho et al., 2004; Nowak et al., 2003; Csicsvari et al., 2003, 1999).

Narrow waveforms, on the other hand, are commonly used as a marker of fast-spiking interneurons, which has been validated with several techniques (Senzai et al., 2019; Mitchell et al., 2007; Gentet et al., 2012; Vinck et al., 2013a; Csicsvari et al., 2003; Hasenstaub et al., 2005; Nowak et al., 2003; Perrenoud et al., 2016; Miri et al., 2018; Vinck et al., 2016; Senzai et al., 2019; Cardin et al., 2009). In many cortical areas, the percentage of NW neurons is comparable to the percentage of GABAergic interneurons established by anatomical studies. For example, in rodent V1 and S1, the percentage of NW neurons is about 10-15% (Vinck et al., 2015; Senzai et al., 2019; Vinck et al., 2016; Rudy et al., 2011; Sirota et al., 2008). Yet, in contrast to previous studies of other brain areas and species, we observed a high (≈70%) percentage of NW neurons in macaque V1. This is in line with Gur et al. (1999), who observed this phenomenon throughout all layers of macaque V1. This percentage is much larger than the percentage of GABAergic interneurons established by anatomical studies (≈20-25%, Hendry et al. (1987)). By analyzing firing properties, we demonstrated that NW neurons formed two distinct classes: NW–Burst and NW–Nonburst neurons.

The firing properties of NW–Nonburst neurons were comparable to those of fast-spiking inhibitory interneurons: 1) NW–Nonburst neurons had late peaks in the autocorrelogram, which indicates a low burst-propensity. Similar firing characteristics have been described for fast-spiking interneurons in other cortical areas (Nowak et al., 2003; Csicsvari et al., 2003; Vinck et al., 2016, 2013a; Senzai et al., 2019; Perrenoud et al., 2016; Csicsvari et al., 1999) (see Figure S2). 2) They had high discharge rates, which is a common characteristic of fast-spiking interneurons (Csicsvari et al., 2003; Nowak et al., 2003; Senzai et al., 2019; Vinck et al., 2013a; Perrenoud et al., 2016) (see Figure S2). 3) They had low stimulus-selectivity, in line with previous studies (Kerlin et al., 2010; Nowak et al., 2007; Perrenoud et al., 2016). 4) They fired with a short time-delay (≈2ms) after neurons in their vicinity (Figure 6D). This is consistent with many models of excitatory-inhibitory interactions, e.g. PING models (Buzsáki and Wang, 2012; Wang, 2010) and previous experimental characterizations (Csicsvari et al., 2003; Hasenstaub et al., 2005; Vinck et al., 2013a).

NW–Burst neurons had early peaks in the autocorrelogram and a high burst-propensity. This high burst-propensity strongly suggests that these neurons are excitatory (Nowak et al., 2003; Csicsvari et al., 2003; Vinck et al., 2016; Senzai et al., 2019; Csicsvari et al., 1999). Which class of excitatory neurons would these NW–Burst neurons correspond to? Based on intracellular recordings and current injections, Gray and McCormick (1996) discovered a subset of NW excitatory neurons in area V1 of the anesthesized cat. These neurons have also been identified in motor and suprasylvian association areas of anesthesized and awake cats (Steriade et al., 1998, 2001). Gray and McCormick (1996) named these neurons “chattering cells” and they resemble the NW–Burst neurons described here in several respects: 1) NW–Burst neurons fired in patterns of high-frequency bursts with an intra-burst frequency up to 400-500Hz, matching the intra-burst frequency of chattering neurons (Nowak et al., 2003; Cardin et al., 2005; Gray and McCormick, 1996; Steriade et al., 1998). Note that intrinsic bursting neurons, which are also excitatory, have broad waveforms and a lower intra-burst frequency (Nowak et al., 2003). 2) NW–Burst neurons had much higher spontaneous discharge rates than BW neurons, consistent with the finding that chattering neurons show little adaptation to depolarizing current injections (Gray and McCormick, 1996; Nowak et al., 2003). By contrast, the responses of intrinsic bursting and regular-spiking neurons adapt strongly over time, and this leads to cycle skipping (Nowak et al., 2003; Gray and McCormick, 1996). Furthermore, in rodent V1 and S1, BW neurons with a high burst-propensity have lower discharge rates as compared to non-bursting BW neurons (Vinck et al., 2015, 2016; Senzai et al., 2019). 3) NW–Burst neurons were strongly stimulus-selective, whereas NW–Nonburst neurons were weakly stimulus-selective. This would be expected if NW–Burst and NW–Nonburst neurons correspond to excitatory and inhibitory neurons, respectively (Kerlin et al., 2010; Nowak et al., 2007; Perrenoud et al., 2016).

However, our data also reveal a major difference in RF properties between NW–Burst and chattering neurons: The majority of chattering neurons in anesthesized cat V1 has simple RFs (Cardin et al., 2005; Nowak et al., 2003; Gray and McCormick, 1996). By contrast, we found that the RFs of NW–Burst neurons were predominantly complex, and that NW–Burst neurons with more complex RFs were more gamma-rhythmic. This discrepancy could reflect a difference between cats and macaques, or a difference between the awake and the anesthesized state.

Our data also suggest that it is unlikely that the sample of NW–Burst neurons included L4 spiny-stellate neurons, which have narrow-waveform spikes (Gur et al., 1999): Whereas spiny-stellate cells are located in L4, our recordings were biased towards superficial layers (L2/3). This is consistent with the presence of strong gamma-synchronization in our dataset, given that gamma is strong in superficial layers and quite weak in L4 (Livingstone, 1996; Xing et al., 2012; Buffalo et al., 2011). In addition, our recorded neurons mainly had complex RFs, which are typical for layers 2/3 but not for layer 4 (Martinez and Alonso, 2003). By contrast, the majority of spiny stellate neurons has simple RF properties (Martinez and Alonso, 2003).

### Unique features of the primate visual cortex

Our findings indicate that primate V1 has unique properties as compared to other areas of the primate cortex and the rodent. In rodent V1, narrow spike-waveforms are a reliable indicator of GABAergic interneurons, and approximately 10-15% of neurons have a narrow waveform (Senzai et al., 2019; Vinck et al., 2015; Rudy et al., 2011) (Figure S2). This also holds true for other cortical areas in the rodent, e.g somatosensory cortex and hippocampus (Vinck et al., 2016; Gentet et al., 2012; Miri et al., 2018). The NW neurons in rodent V1 and S1 do not exhibit burst-firing (Senzai et al., 2019; Vinck et al., 2015, 2016) (Figure S2). In macaque V4 and PFC (prefrontal cortex), the percentage of narrow-waveform neurons is also much lower than in macaque V1 (Vinck et al., 2013a; Mitchell et al., 2007; Ardid et al., 2015). Like the NW–Nonburst neurons in macaque area V1, NW neurons in V4 have high discharge-rates and do not exhibit bursting (Vinck et al., 2013a). As discussed above, intracellular recordings in anesthesized cat V1 (as well as motor and association areas, Steriade et al. (1998)) have identified a subset of excitatory (“chattering”) neurons with narrow waveforms (Gray and McCormick, 1996; Nowak et al., 2003). Previous studies had observed that gamma-rhythmic firing in cat and monkey V1 is often accompanied by burst firing (Livingstone, 1996; Gray et al., 1990; Gray and Di Prisco, 1997; Hubel and Wiesel, 1965; Friedman-Hill et al., 2000). We were surprised that such a large fraction of macaque V1 neurons had firing properties similar to the ones of chattering neurons. Similar firing patterns were observed in a new-world Capuchin monkey (Figure S3A-B). We conclude that monkey V1 has a unique composition of neuron types as compared to rodent V1 and other cortical areas of the monkey. It is unlikely that this is due to the specific behavioral task or state, because NW–Burst neurons also had a high burst-propensity in the pre-stimulus period (Figure 3 and S4). Rather, this difference might reflect expression of specific channels in excitatory neurons like Kv3 voltage-gated potassium channels, which has been observed in macaque V1 but not rodent V1 (Constantinople et al., 2009). Notably, single-cell transcriptional analyses suggest that the properties of excitatory neurons may vary greatly across cortical areas, in contrast to GABAergic interneurons (Tasic et al., 2018). In sum, there are major differences in the waveform and firing properties of excitatory neurons between rodent and macaque V1, and between different areas of macaque cortex.

Another prominent, and possibly related, feature of cat, monkey and human V1 (and V2) is that many visual stimuli induce a prominent 30-80Hz gamma-band rhythm (Gray et al., 1989; Fries, 2009; Vinck and Bosman, 2016; Roberts et al., 2013; Brunet et al., 2014a; van Pelt et al., 2012; Gray and Di Prisco, 1997; Gray et al., 1990) This V1 gamma-rhythm differs in several respects from gamma rhythms in other species and brain areas: (1) Spike-field locking is extremely strong in primate V1 as compared to other brain areas, or rodent V1 (Vinck and Bosman, 2016). In macaque V1, spike-field PPC values range between 0.05-0.15 during visual stimulation with gratings (Figure 6). By comparison, in macaque V4, rodent sensory cortex and hippocampus, gamma PPC values are typically smaller than about 0.005 (Vinck et al., 2013a; Perrenoud et al., 2016; Vinck et al., 2016; Cabral et al., 2014; van Wingerden et al., 2010). (2) In a given individual and stimulus condition, the frequency-range of gamma in cat and primate V1 is quite narrow (about ±10Hz), which is evident from very slowly decaying side-lobes in the autocorrelogram and the spike-triggered-average of the LFP (Figure 5 and 6). By comparison, in rodent S1 and V1, gamma phase-locking occurs in a broad frequency range (30-100Hz) (Vinck et al., 2013a; Perrenoud et al., 2016; Vinck et al., 2016) (but see Veit et al. (2017), who reported a peak in the LFP around 25-30Hz in mouse V1). This is reflected in spike-triggered LFP averages and autocorrelograms that have rapidly decaying side-lobes (Perrenoud et al., 2016; Vinck et al., 2016). (3) Many visual stimuli cause a dramatic increase in cat and monkey V1 gamma-band power relative to pre-stimulus baseline, which can be as great as 10 to 300-fold (Brunet et al., 2014b; Peter et al., 2019; Gray et al., 1989, 1990; Gray and Di Prisco, 1997; Gieselmann and Thiele, 2008; Lima et al., 2010). By contrast, in mouse V1, grating-stimuli increase LFP gamma-power only by about 1.2-to 2-fold relative to baseline (Vinck et al., 2015; Perrenoud et al., 2016; Veit et al., 2017).

Given their extremely strong phase-locking, we expect that NW–Burst neurons play a crucial role in the generation of V1 gamma oscillations. This would mean that the gamma-rhythm in primate and cat V1 depends on specialized mechanisms that may not be present in many other brain areas and species. Why would gamma in macaque V1 (and perhaps extrastriate visual areas) have these unique features? We have previously proposed that in the visual system, gamma-synchronization reflects predictability among visual inputs across space (Vinck and Bosman, 2016; Peter et al., 2019). This hypothesis predicts high-amplitude gamma when RF sizes are small, because in natural scenes, predictability between visual inputs should decrease as a function of distance (Vinck and Bosman, 2016). Thus, when the RFs are small as in case of macaque V1, bottom-up visual inputs should be more predictable from the spatial surround (Vinck and Bosman, 2016). A complementary explanation is that when cortical networks are large as is the case in cat and monkey V1 (large magnification factor and small RFs), reciprocal communication and cooperation between distal nodes may require strong oscillatory synchronization (Singer, 2018).

### Mechanistic and functional implications

Gamma oscillations in the 40-90Hz range are thought to depend on interactions between excitatory neurons and fast-spiking interneurons (Wang, 2010; Buzsáki and Wang, 2012; Cardin et al., 2009; Hasenstaub et al., 2005; Chen et al., 2017; Kopell et al., 2000; Perrenoud et al., 2016; Vinck et al., 2013b; Börgers and Kopell, 2008, 2005; Lytton and Sejnowski, 1991; Jadi and Sejnowski, 2014; Whittington et al., 1995; Sohal et al., 2009; Wang and Buzsáki, 1996; Womelsdorf et al., 2014b; Bush and Sejnowski, 1996; Whittington et al., 2011, 2000; Bartos et al., 2007). Experimental studies have found that fast-spiking interneurons are more gamma phase-locked than (BW) excitatory neurons (Senzai et al., 2019; Vinck et al., 2013a; Csicsvari et al., 2003; Hasenstaub et al., 2005; Perrenoud et al., 2016; Salkoff et al., 2015; Vinck et al., 2016; Schomburg et al., 2014; Bragin et al., 1995). In macaque V1, we found that putative fast-spiking interneurons (NW–Nonburst neurons) had much lower gamma phase-locking than NW–Burst neurons (Figure 6). In fact, their precision of phase-locking was similar to BW neurons, although a few NW–Nonburst neurons had rhythmic side-lobes in the autocorrelogram (Figure 2). These results are consistent with data from intracellular recordings in anesthesized cat V1 (Azouz et al., 1997), in which the autocorrelograms of most fast-spiking interneurons did not have prominent rhythmic side-lobes. On the other hand, we found signatures of strong rhythmic inhibition in NW–Burst neurons: First, we found that the discharge rates of NW–Burst neurons were suppressed below pre-stimulus levels during their non-preferred gamma phase (Figure 7). This indicates an inhibitory input arriving at their non-preferred gamma phase. Second, we found that the discharge rates of most NW–Burst neurons were suppressed at the non-preferred stimulus orientations. This suggests that they receive a broadly-tuned inhibitory drive, which may sharpen their orientation tuning (Shapley et al., 2003; Kerlin et al., 2010; Priebe and Ferster, 2008; McLaughlin et al., 2000; Isaacson and Scanziani, 2011). Yet, despite this firing suppression for suboptimal stimuli, the gamma phase-locking of NW–Burst neurons remained strong (Figure 8), indicating entrainment by rhythmic inhibition. If both orientation tuning and gamma phase-locking depend on inhibitory drive, then this might also explain why the most gamma phase-locked NW–Burst neurons were the most orientation-tuned (Figure 8).

Although gamma-synchronization likely reflects a generic dynamical motif emerging from recurrent E-I interactions, the strength and frequency-bandwidth of gamma could be highly dependent on the specific properties of excitatory neurons. These properties may vary across cortical areas, species or laminae (Tasic et al., 2018; Senzai et al., 2019). In most (and perhaps all rodent) cortical systems, fast-spiking interneurons interact with broad-waveform excitatory neurons that are regular spiking or irregularly bursting, and exhibit low-pass filtering characteristics (Hasenstaub et al., 2005; Pike et al., 2000; Cardin et al., 2009). Fast-spiking interneurons on the other hand have a subtreshold transfer function that shows less low-pass filtering, and have a suprathreshold transfer function that is relatively flat across frequencies up to 100Hz (Hasenstaub et al., 2005). Experimental studies have found that excitatory neurons with broad waveforms are weakly phase-locked compared to fast-spiking interneurons, and fire in only a small percentage of gamma cycles (i.e. have low discharge-rates and skip cycles) (Hasenstaub et al., 2005; Csicsvari et al., 2003; Vinck et al., 2013a, 2016; Perrenoud et al., 2016; Senzai et al., 2019; Schomburg et al., 2014; Vinck et al., 2015). In fact, fast-spiking interneurons in mouse V1 and macaque V4 can sustain gamma-band oscillations despite very weak or no entrainment of excitatory neurons, e.g. in the pre-stimulus baseline period (Perrenoud et al., 2016; Vinck et al., 2013a; Batista-Brito et al., 2017).

On the other hand, in macaque V1, the nature of E-I interactions appears to be fundamentally different. We found that NW–Burst neurons exhibited very strong gamma-phase locking and high discharge rates. This suggests that NW–Burst neurons integrate synaptic inputs as a band-pass filter, exhibiting resonance in the gamma-frequency band. They would thereby transform rhythmic inhibitory inputs into highly rhythmic spiking-outputs, promoting high-amplitude, narrow-band gamma oscillations in the network. In support of this resonance or pacemaker hypothesis, a recent study in anesthesized cats found that optogenetic stimulation of excitatory neurons induced the strongest change in oscillatory power and phase-locking when the stimulation frequency was in the gamma-frequency band (Ni et al., 2017). By contrast, in rodent S1 and prefontal cortex, excitatory neurons show characteristic low-pass filtering (Cardin et al., 2009; Hasenstaub et al., 2005), consistent with the idea that cat and primate V1 have unique features. In sum, when fast-spiking interneurons interact with broad-waveform excitatory neurons, the resulting gamma-rhythm occurs in a relatively broad frequency-band and is comparably weak, with stronger phase-locking in interneurons than in BW excitatory neurons. However, when fast-spiking interneurons interact with NW, bursting excitatory neurons that have resonance in the gamma range, the resulting gamma rhythm can be very strong and occur in a narrow frequency range, with very strong gamma phase-locking in NW, bursting excitatory neurons. As an extension to the standard PING model, we call this gamma-model PRING (pyramidal-resonance interneuron network gamma).

The association of bursting with gamma-phase locking raises the question whether these two processes are causally related (Womelsdorf et al., 2014a). We found that the burst propensity of NW–Burst neurons was similar in the visual-stimulation and pre-stimulus periods, i.e. in periods with strong and weak to absent gamma, respectively. This indicates that their burst firing is neither a consequence of visual stimulation, nor of visually-induced gamma synchronization, and that burst firing does not always lead to network gamma. Rather, our data suggests that bursts occur asynchronously in the pre-stimulus period, and become strongly synchronized across neurons during visual stimulation.

The properties of NW–Burst neurons suggest that they have a particularly strong impact on post-synaptic targets: 1) Burst-firing can increase the gain and reliability of synaptic transmission, and facilitate long-term potentiation (Lisman, 1997; Jackman and Regehr, 2017). 2) Synchronization of spikes can increase the gain of transmission to post-synaptic targets (Salinas and Sejnowski, 2000; Fries, 2015; Vinck et al., 2013b; Knoblich et al., 2010; Abeles, 1982; Palmigiano et al., 2017; König et al., 1996; Bernander et al., 1994; Kempter et al., 1998; Womelsdorf et al., 2007; Ni et al., 2016) and feedforward communication between visual areas is especially strong at gamma-frequencies (Bastos et al., 2015; van Kerkoerle et al., 2014; Michalareas et al., 2016; Buschman and Miller, 2007; Gregoriou et al., 2009). Gamma-synchronization is also associated with fast behavioral reaction-times (Rohenkohl et al., 2018; Siegle et al., 2014; Womelsdorf et al., 2006). Further, gamma-synchronization may have important consequences for spike-time-dependent-plasticity (Vinck et al., 2010a; Sejnowski and Paulsen, 2006; Markram et al., 1997; Galuske, ????). Together with their high stimulus-selectivity, this suggests that NW–Burst neurons could be the principal source of communication with downstream target areas.

In sum, macaque V1 contains a specialized cell-type that fires in high-frequency bursts, is strongly phase-locked to gamma oscillations, and is highly stimulus-selective. This neuron type is likely pivotal for the encoding and transmission of stimulus information. Through interactions with inhibitory neurons, these neurons could act as pacemaker for V1 gamma oscillations.

## Methods

### Ethics

All procedures complied with the guidelines of the European Community for the care and use of laboratory animals (European Union Directive 86/609/EEC). The experiments were approved by the regional authority for animal welfare (Regierungspräsidium Hessen, Darmstadt, Germany).

### Surgical procedures

Recordings were made from two adult rhesus-monkeys (Macaca mulatta). We performed surgical procedures under general anesthesia and provided analgesic treatment after the operations. In each monkey, we implanted a titanium headpost, which was later used for head fixation, and a titanium recording chamber. We inserted 2 to 10 microelectrodes independently into the cortex via transdural guide tubes (diameter: 300 *μ*m; Ehrhardt Soehne). These electrodes had distances to each other between 1 and 3mm and were made of quartz-insulated, tungstenplatinum material (diameter: 80*μ*m; impedances between 0.3 and 1MΩ; Thomas Recording). We recorded from superficial layers in the opercular region of V1. The receptive-field centers of the units had eccentricies between 2 and 3 degrees of visual angle.

### Recordings

Electrode signals were amplified (1000x) and filtered using a 32-channel head-stage amplifier (head stage HST16o25; head stage and preamplifier from Plexon). The filter-ranges for multi-unit activity and LFPs (Pesaran et al., 2018) were 0.7-6kHz and 0.7-170Hz, respectively. An on-board acquisition board provided an additional 10x amplification (E-series acquisition boards; National Instruments). The signals were digitized and stored with a LabVIEW-based acquisition system (written by SN). To detect spikes, we manually set a threshold based on online visualization of the spike waveforms. This threshold was typically between 2 and 3 SDs above the noise level. LFP signals and spike waveforms were sampled at 1kHz and 32kHz, respectively. The eye position was monitored either with a search coil system (DNI, Crist Instruments, USA; temporal resolution of 2 ms) or with an infrared eye tracker (Matsuda et al. 2000; temporal resolution of 33 ms).

### Behavioral training and task

We trained both monkeys on a fixation task. At the start of each trial, a square, red fixation-dot appeared on the monitor (0.15 degrees; 4 × 4 pixels; luminance: 10.0 cd/m^2^). The monkeys then had to press a lever, and maintain their gaze within a radius of ≈0.5 degrees of visual angle from the center of the fixation point. After a random interval within 2500-4000ms, the color of the fixation dot changed from red to green. The monkey could then obtain a reward by releasing the lever between 200 and 500ms after the color change. A trial was aborted if, before the color change, the monkey released the lever or moved his eyes outside the fixation window. The monkeys typically performed between 700 and 1500 correct trials in a 4-h session.

### Visual stimulation paradigm during recordings

We used an interface in LabVIEW (written by SN; Lab-VIEW, National Instruments, USA) to generate stimuli as sequences of bitmap images. The stimuli were presented as movies with a resolution of 1024 × 768 pixels and a frame rate of 100-120Hz. We controlled the stimulus presentation with a graphical board (GeForce 6600-series, NVIDIA, Santa Clara, CA) and the ActiveStim software (www.activestim.com). With the ActiveStim software, we achieved accurate timing and a stimulus-onset jitter below 1ms. The movies were displayed on a cathode-ray-tube monitor (CM813ET, Hitachi, Japan), which was 36 × 28 degrees of visual angle wide (1024 × 768 pixels). We gamma-corrected this monitor, such that the relationship between output luminance and gray values was linear.

At the beginning of each recording session, we presented moving bar-stimuli in 16 different directions. We analyzed the neural responses to these bar stimuli in order to estimate the RFs of the recorded units. We then proceeded with the presentation of drifting-grating stimuli, which were centered on the RFs. Each trial started with a pre-stimulus baseline of 800-1000ms, followed by the presentation of a drifting-grating stimulus for a duration of 800-1500ms. The stimulus duration was constant in a given recording session. The drifting gratings had spatial frequencies of 1.25-2 cycles per degree, and temporal frequencies of 1.4-2 cycles per second. In each trial, the direction of the grating drift was orthogonal to grating orientation and randomly chosen from 16 directions (in steps of 22.5 degrees). During the presentation of visual stimuli, the monkeys performed a fixation task, which means that the stimuli had no behavioral relevance for the monkeys.

### Recordings in Capuchin monkeys

Additional data was obtained from recordings in one capuchin monkey (Sapajus libidinosus). All procedures related to recordings in capuchin monkeys were approved by the Ethics Committee of the Federal University of Rio Grande do Norte (Protocol number 053/2012, CEUA). Capuchins were required to maintain fixation continuously for 2000 to 3000 ms. Different from the experiments with the macaque monkeys, the capuchins were not required to press a lever. All other procedures (electrodes, recording devices, acquisition and stimulation systems) were identical between the two experiments.

### Recordings in mice

The details for recordings in mice are described in detail in (Hoy and Niell, 2015). We recorded from area V1 in awake, adult head-fixed C57BL/6J mice placed on a spherical treadmill (Niell and Stryker, 2010). Recordings were made with two-shank 32-channel silicon probes having 25*μ*m spacing between channels (Neuronexus). Electrodes were coated in DiO, which we used for identification of the laminar position of the electrodes. Data acquisition was performed using the TDT System 3 workstation. We presented full-field drifting-grating stimuli with spatial frequencies between 0.01 and 0.32 (0.01, 0.02, 0.04, 0.08, 0.16, 0.32) and 16 different drift-directions. The extracellular signal was filtered between 0.7 to 7 kHz and stored it at 25 kHz. We detected spiking events online by voltage threshold crossings. For each spike, we stored a 1 ms waveform sample on four adjacent recording surfaces, which formed a “virtual” tetrode that was then used for spike sorting. We performed single-unit clustering and spike-waveform analysis as described in Niell and Stryker (2010); Hoy and Niell (2015).

### Spike-sorting

We performed semi-automated spike sorting to isolate neurons. For automatic clustering, we used the KlustaKwik 3.0 software. The features used were the energy of the spike waveform, and the energy of the waveform’s first derivative. Based on the results of the automatic clustering, we manually sorted units with the M-Clust software. We accepted units only if a clear separation of the cell relative to the noise clusters was observed, and if the inter-spike-interval distribution had a clear refractory-period. This was generally the case when a conventional measure of cluster separation, isolation distance (ID) exceeded 20 (Schmitzer-Torbert et al., 2005) (the median ID was 25.05). NW–Burst and NW–Nonburst neurons did not significantly differ in the ID (*P* > 0.05).

### Data analysis

We analyzed the data in MATLAB by using the FieldTrip toolbox (Oostenveld et al., 2011) and custom-made scripts (IO and MV). We included only correct trials into our analyses.

### Computation of the autocorrelogram

We computed the autocorrelogram separately for the stimulus period (>200ms after stimulus onset) and the pre-stimulus period. The autocorrelogram was computed by sampling the spike trains at 2000Hz and convolving them with themselves. This corresponds to counting the number of times that a pair of spikes has a certain delay. We corrected the autocorrelogram for bias due to finite analysis periods. We estimated the time at which the autocorrelogram reached a global peak with the following procedure:

1. Because the autocorrelogram sampled at 2000Hz can have high variability across samples, we fitted a polynomial to the autocorrelogram. We determined the order of the polynomial fit with a cross-validation procedure, which prevented overfitting. In this cross-validation procedure, trials were divided into two non-overlapping sets, which were the “training” and the “test” set. Polynomials of orders from 5 to 40 were fitted on the training set. We selected the polynomial order that yielded the smallest error on the test set. Finally, we fitted a polynomial of the selected order to the autocorrelogram that was computed based on all the trials.
2. Because the autocorrelograms often had sidelobes due to rhythmic firing, we dampened this rhythmicity by convolving the polynomial fits with a moving-average filter. The length of the moving-average filter was equal to the average length of the gamma cycle, which was determined per monkey separately (15ms in monkey J and 26ms in monkey L). The moving-average filter was only applied to samples beyond 8ms.
3. We determined the time at which the smoothed polynomial-fit reached a global peak within 0-60ms (Figure 2E). In addition, we determined the time at which the smoothed autocorrelogram reached a global peak within 0-12ms (Figure S1C).
4. We defined two additional measures of burst-propensity. The first measure of burst-propensity was comparable to a Z-score, and was defined as 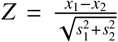. Here, *x*_1_ was the maximum value of the polynomial fit within 5ms, and *x*_2_ was the minimum value within 5-10ms. These values were associated with standard errors *s*_1_ and *s*_2_, which were obtained through a bootstrapping procedure. The value of this measure is shown in Figure 2E. We defined another measure of burst-propensity by comparing the maximum value of the polynomial fit within 0-5ms, *x*_1_, with the maximum value within 8-10ms, *x*_2_. This measure was defined as 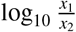 and is shown in Figure S1B-C.
5. We identified two non-overlapping clusters of NW neurons based on the global peak-time in the autocorrelogram (Figure 2E). As an additional criterion for NW–Burst neurons, we required them to have a burst-modulation of at least 20%. This criterion was not reached by two neurons (Figure S1B), which were not further analyzed. Inclusion of these two neurons into the NW–Burst population did not qualitatively alter the results. A few neurons had peaks in the autocorrelogram within 5ms, and a relatively high burst-modulation score (Figure S1B). However, their autocorrelograms did not reach a global maximum within 5ms when considering the entire 60ms interval (Figure S1C). We considered the classification of these neurons as unclear and included them neither into the NW–Burst nor in the NW–Nonburst population.

### Other measurement of burst-propensity

The total fraction of burst-spikes was computed based on the inter-trial-interval distribution. We counted the number of spikes either between 0.008ms and 3.5ms (short burst intervals), or between 0.008ms and 6ms (long burst intervals). We defined the ratio *a*/*b*, where *a* was the number of burst spikes and *b* the total number of spikes.

To compute the correlation between burstiness in the prestimulus and the stimulus period (Figure S4), we defined the burst-propensity based on the autocorrelogram. This avoids potential biases due to differences in discharge rates (Senzai et al., 2019). The burst-propensity was defined as the ratio (*a* − *b*)/(*a* + *b*). Here, *a* was the average value of the normalized autocorrelogram either between 0.0008s and 3.5ms (short bursts), or between 0.0008s and 6ms (long bursts). The variable *b* was the average value of the normalized autocorrelogram between 0.008s and 0.1s. We also used this measure of burst-propensity for the correlation with spike-LFP phase-locking values.

### Quantification of spike-LFP phase-locking

To compute spike-LFP phase-locking, we determined the phase of each spike relative to the LFP recorded from the other electrodes. To remove line noise, we filtered the LFP signals with a two-pass 4^th^ order Butterworth bandstop-filter (49.5-50.5Hz). We then averaged the LFP signals across the other electrodes, which was justified because of their high coherence. We computed the “spike-LFP phase” in two different ways:

In the first approach, we computed the wavelet transform of the LFP snippet around each spike, using a constant number of cycles (9) per frequency (as in (Vinck et al., 2013a)). The advantage of this method is that it allows for a comparison of spike-LFP phase-locking between different frequencies, similar to the spike-field coherence. The disadvantage is that many LFP cycles are used to determine the spike-LFP phase. This could lead to an underestimation of spike-LFP phase-locking, because the probability of a spike should not be influenced by LFP fluctuations that are far away in time.

The second method addressed this problem and consisted of the following steps:

1. We band-pass-filtered the LFP in a relatively broad frequency-band with a 3-th order, two-pass Butterworth filter (40-90Hz in monkey J, 25-55Hz in monkey L).
2. We computed the Hilbert-transform of the band-pass filtered signal and determined the instantaneous phase.
3. We detected gamma cycles as follows: First, we detected all the zero-crossings of the instantaneous phase, which occur in the neighbourhood of peaks in the band-pass filtered signal. For each k-th zero-crossing, we examined whether the angular velocity of the phase was positive for all time points between the *k* − 1-th to the *k* + 1-th zero-crossing. If this was not the case, then there was a “phase-slip” in which the instantaneous frequency became negative. Otherwise, we detected the nearest peak in the bandpass filtered signal relative to the *k*-th zero-crossing of the phase. We then measured the duration of the gamma cycle as the time from the current peak to the next peak in the signal.
4. We ran the same cycle selection-procedure on the prestimulus period, in which narrow-band gamma-band oscillations are virtually absent. For the pre-stimulus period, we obtained the mean *μ_pre_* and standard deviation *σ_pre_* of the distribution of amplitudes. These amplitudes were measured as the peak-to-valley distance of the gamma cycle (Atallah and Scanziani, 2009). A cycle in the stimulus period with amplitude *A* was only selected if (*A* − *μ_pre_*)/*σ_pre_* > 1.63 (which would be the cut-off for a one-sided T-test at *P* < 0.05).
5. We defined the gamma phase as *t*/*T*, where *t* was the time from the peak of the gamma cycle, and *T* the time to the next peak.

After computing the spike-LFP phases, we estimated phase-locking with the pairwise-phase-consistency (PPC) (Vinck et al., 2012, 2010b). Specifically, we used the PPC1 measure defined by Vinck et al. (2012). The PPC1 takes all pairs of spikes-LFP phases from separate trials and computes the average consistency of phases across these pairs. It is not affected by mean discharge-rates and history effects like bursting (see Methods) (Vinck et al., 2012). The bias due to discharge rates is removed by the pairwise computation (Vinck et al., 2012). The bias due to history effects is removed by considering only pairs of spike-LFP phases from separate trials (Vinck et al., 2012). Thus, PPC values can be directly compared between neuron types with varying discharge rates and burst-propensity. Because phase-locking estimates can have a high variance for low spike counts, we computed PPC values only for neurons that fired at least 50 spikes (Vinck et al., 2013a).

To compute spike-triggered averages of the LFP, we first Z-transformed the LFP signals such that they had unit standard deviation. The STA was defined as the average Z-scored LFP signal across spikes.

### Orientation and direction tuning

We measured orientation-tuning with the orientation-selectivity-index (OSI). The OSI was defined as in Womelsdorf et al. (2012): The mean discharge rates, *r_m_* (spikes/sec), were determined for each *m*-th stimulus orientation. The angle of the stimulus orientation *θ_m_* was a circular variable that ranged between 0 and 2*π* radians. The discharge rates were normalized as 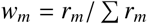. The OSI was defined as

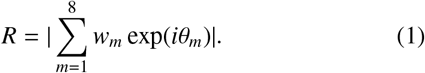

This quantity can be understood as the resultant vector-length, which is obtained by adding eight vectors with angle *θ_m_* and magnitude *w_m_*. For the direction-selectivity index (DSI), the variable *θ_m_* represented the stimulus direction and *w_m_* was the discharge rate for the *m*-th stimulus direction. We have previously shown that the OSI estimate is positively biased (Womelsdorf et al., 2012). This bias decreases as the discharge rate or the trial count increase (Womelsdorf et al., 2012). Thus, uncorrected OSI (and DSI) values are typically high for neurons with low discharge-rates (Womelsdorf et al., 2012). We estimated this bias by shuffling the discharge rates across trials, and computing *R* for the shuffled data (Batista-Brito et al., 2017). We then computed the average value of *R* across 1000 shuffling iterations, and subtracted the bias estimate from the observed value of the OSI (or DSI) (Batista-Brito et al., 2017).

### Simple vs. complex-cell modulation

We used the following procedure to determine the extent to which V1 neurons had simple or complex receptive-fields: For each neuron, we estimated the discharge rate as a function of time by convolving the spike train with a Gaussian kernel (40ms length, standard deviation 10ms). “spike-density function”). This spike-density function was computed only for the preferred stimulus-orientation. We then fitted a sinusoid with the temporal frequency of the drifting gratings to this spike-density function. We computed this fit for the period starting > 200ms after stimulus onset, in order to avoid the initial stimulus transient. We also estimated the bias of the sinusoid fit, *B*, by fitting a sinusoid which was shifted by a random phase in each trial. A modulation term *F*1 was defined as the difference between the amplitude *A* and the bias *B, F*1 = *A* − *B*. If *F*1 < 0, then we set *F*1 = 0. To measure simple vs. complex-cell modulation, we computed the measure *M* = (*F*1 − *F*0)/(*F*0 + *F*1). Here, *F*0 is the mean elevation of firing above baseline. Note that this definition is different from the F1/F0 ratio that has been used in other studies (Martinez and Alonso, 2003). However, the normalization in the denominator makes *M* robust against outliers and also well-behaved when *F*0 is close to zero. Previous studies have used *F*1/*F*0 = 1 as a cut-off point to define simple cells. With the definition of *M*, the equivalent cut-off point for simple cells would be *M* = 0.

### Correlation between firing rate and phase-locking across stimuli

We computed the correlation between firing rate and phase-locking across the 16 stimulus directions as follows (Figure 8): For each neuron, we determined discharge rates for each of the 16 different stimulus-directions. These discharge rates were normalized by dividing by the maximum discharge-rate, such that the normalized discharge-rates were bound by 0 and 1. We also computed the gamma phase-locking value (PPC) for each of the 16 stimulus directions separately. The 16 gamma phase-locking values (PPC) were then predicted from the 16 corresponding values of the discharge rate. Because the variance of the phase-locking estimate (PPC) decreases as a function of the spike count, we used a weighted linear-regression model. In this model, the influence of an observation on the regression fit was proportional to the inverse of the variance. The variance of the PPC was estimated with a bootstrapping procedure, in which we bootstrapped over trials.

### Statistics

We used permutation testing for statistical comparisons between different neuron types (Maris and Oostenveld, 2007). First note that the two monkeys may systematically differ in a certain variable (e.g. the OSI), and that there might be more neurons of a certain type in one monkey. This could potentially bias our analyses when comparing neuron types. We addressed this problem as follows: For a comparison between values of *X* (representing any variable) between two neuron types, we Z-scored the values of *X* across the two neuron types per monkey separately. This ensured that the mean of the Z-scored variable was identical between the two monkeys. We then pooled the Z-scores across the two monkeys. As an alternative testing strategy, we also computed the difference in *X* per monkey separately, and averaged the difference score across the two monkeys. This yielded qualitatively similar results. For permutation testing, we randomly permuted the labels of the two neuron types under consideration. We then constructed a permutation distribution for differences between two neuron types. We computed the P-value by comparing the observed difference between two neuron types with the permutation distribution. We corrected for multiple comparisons using an FDR correction (5% false positive rate, correction for dependent tests). Error bars in all figures correspond to standard errors of the mean.

---

**Figure S1:**
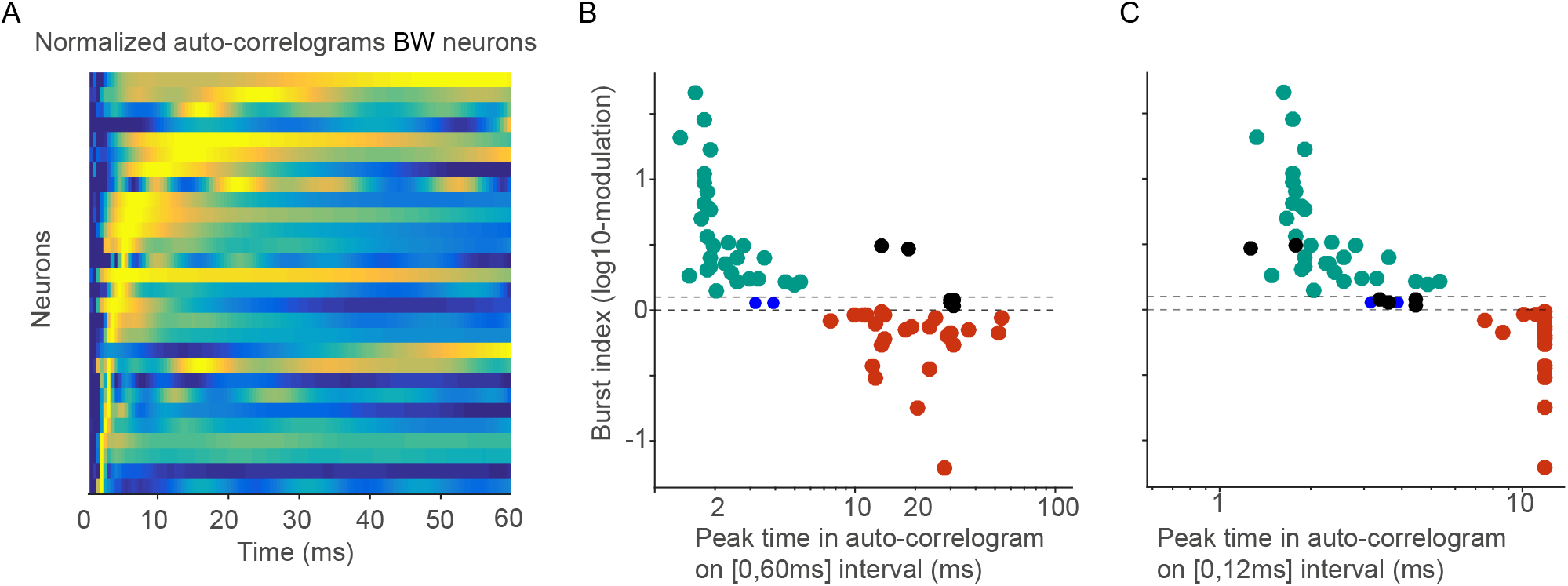
(A) Similar to Figure 2C-D, but now for BW neurons. (B) The time at which the autocorrelogram reached a global peak within 60ms (x-axis) vs. burstiness. Burstiness was measured as the log10-ratio of the autocorrelogram maximum within 5ms vs. the maximum at 8-10ms (see Methods). (C) Similar to (B), but now the time at which the autocorrelogram reached a global peak within 12ms. Note that a few neurons reached an early peak in the autocorrelogram when considering the 12ms interval, but not the 60ms interval (in black). These neurons also had higher levels of burstiness. We excluded these neurons from further analyses, i.e. did not assign them to either the NW–Bursting or NW-Non-bursting class. Furthermore, there were two neurons that had very low levels of burstiness (< 0.1), close to the levels of burstiness in the NW-Non-bursting neurons (in blue). We excluded these neurons from further analyses, i.e. did not assign them to either the NW–Bursting or NW-Non-bursting class. Results did not qualitatively change with inclusion of these neurons, however.

**Figure S2:**
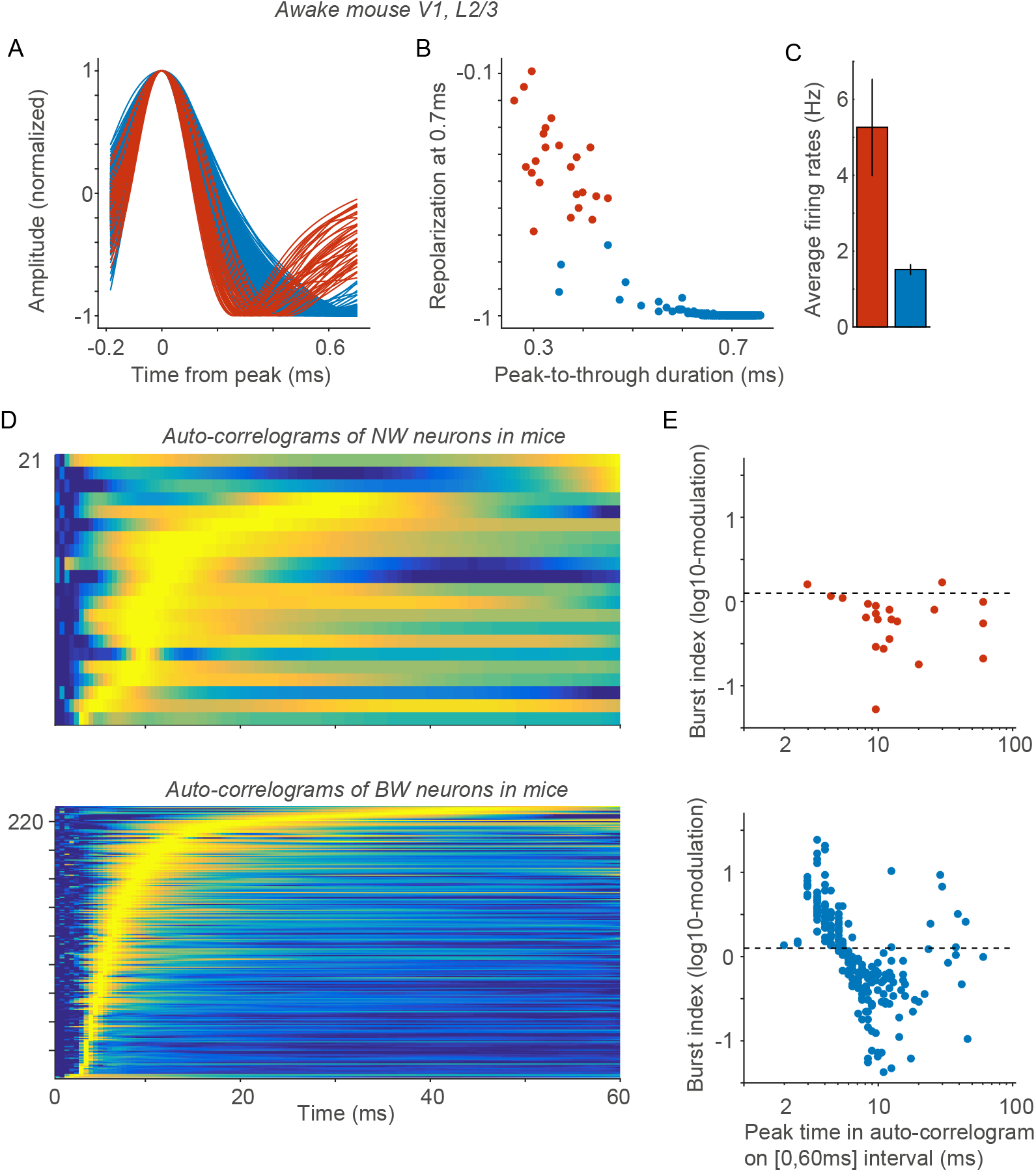
(A) Action potentials waveform for L2/3 neurons in mouse V1. (B) AP peak-to-trough duration vs. repolarization at 0.7 ms. NW neurons were defined as having repolarization values larger than −0.7. (C) Spontaneous discharge rates (spikes/sec). (D) Autocorrelograms for NW and BW neurons in L2/3 of mouse V1. These autocorrelograms were computed for periods of visual stimulation with drifting gratings (> 0.2s after stimulus onset). This figure is similar to Figure 2C-D. The only difference is that we did not smooth the autocorrelograms, because there is no strong gamma-rhythmicity in this data as in case of macaque V1. (E) The time at which the autocorrelogram reached a global peak within 60ms (x-axis) vs. burstiness, separate for NW and BW neurons in L2/3 mouse V1 neurons. Burstiness was defined as in Figure S1B. This figure is similar to Figure S1B. Only one of the NW neurons passed our criteria to be a NW–Burst neuron. Note that many of the BW neurons present burst-firing. Very few neurons however reach a peak in the autocoπelogram below 3ms, which is a main difference to the NW–Burst neurons in macaque V1. This means that NW–Burst neurons in macaque V1 have higher intra-burst frequencies than BW neurons in mouse V1.

**Figure S3:**
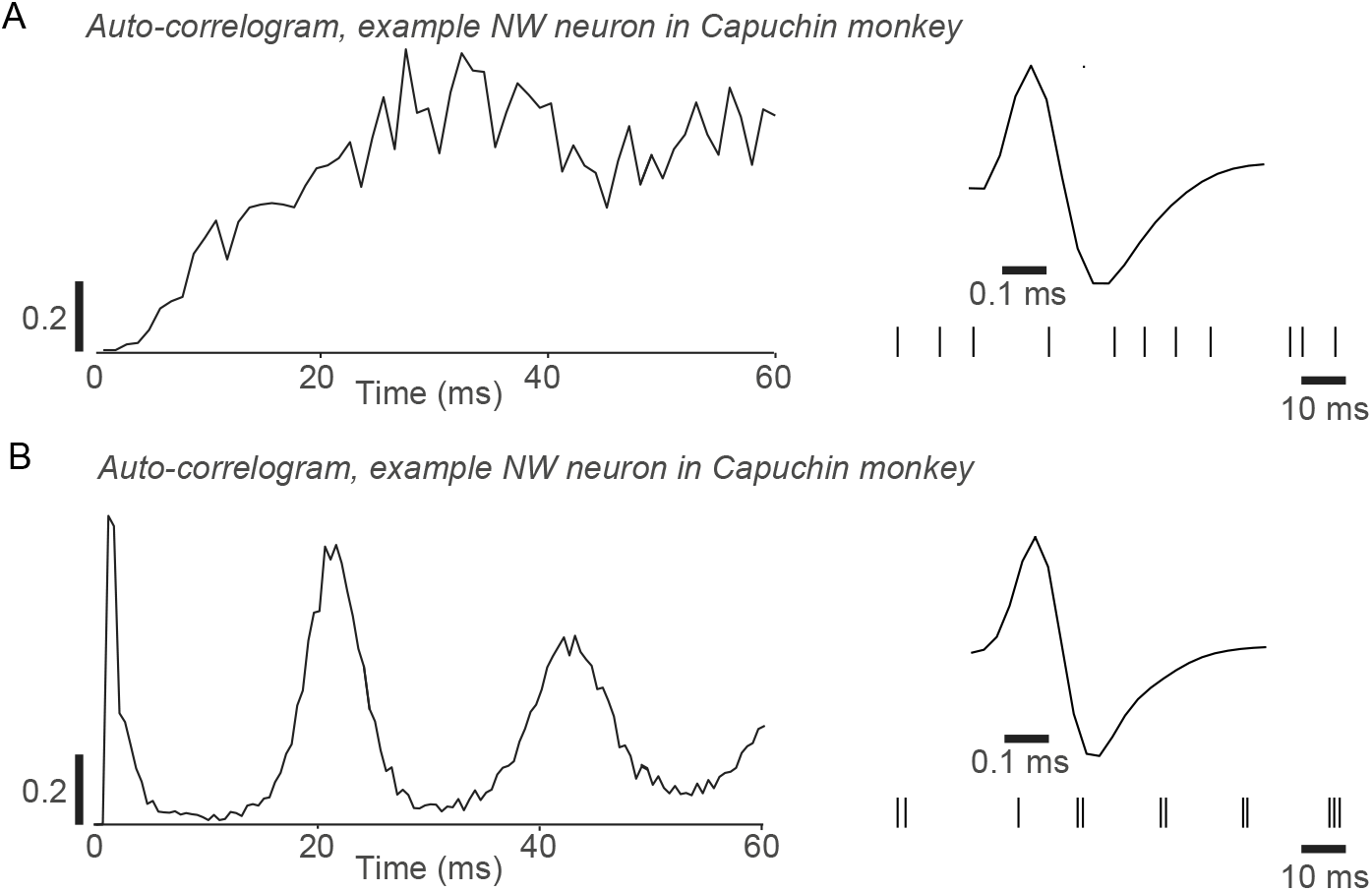
(A) Example NW neuron that was recorded from area V1 of the awake Capuchin monkey. Recordings were made during presentation of drifting-grating stimuli. The firing behavior of this neuron is similar to the one of NW–Nonburst neurons. (B) Example NW neuron that was recorded from area V1 of the awake Capuchin monkey. The firing behavior of this neuron is similar to the one of NW–Bursting neurons.

**Figure S4:**
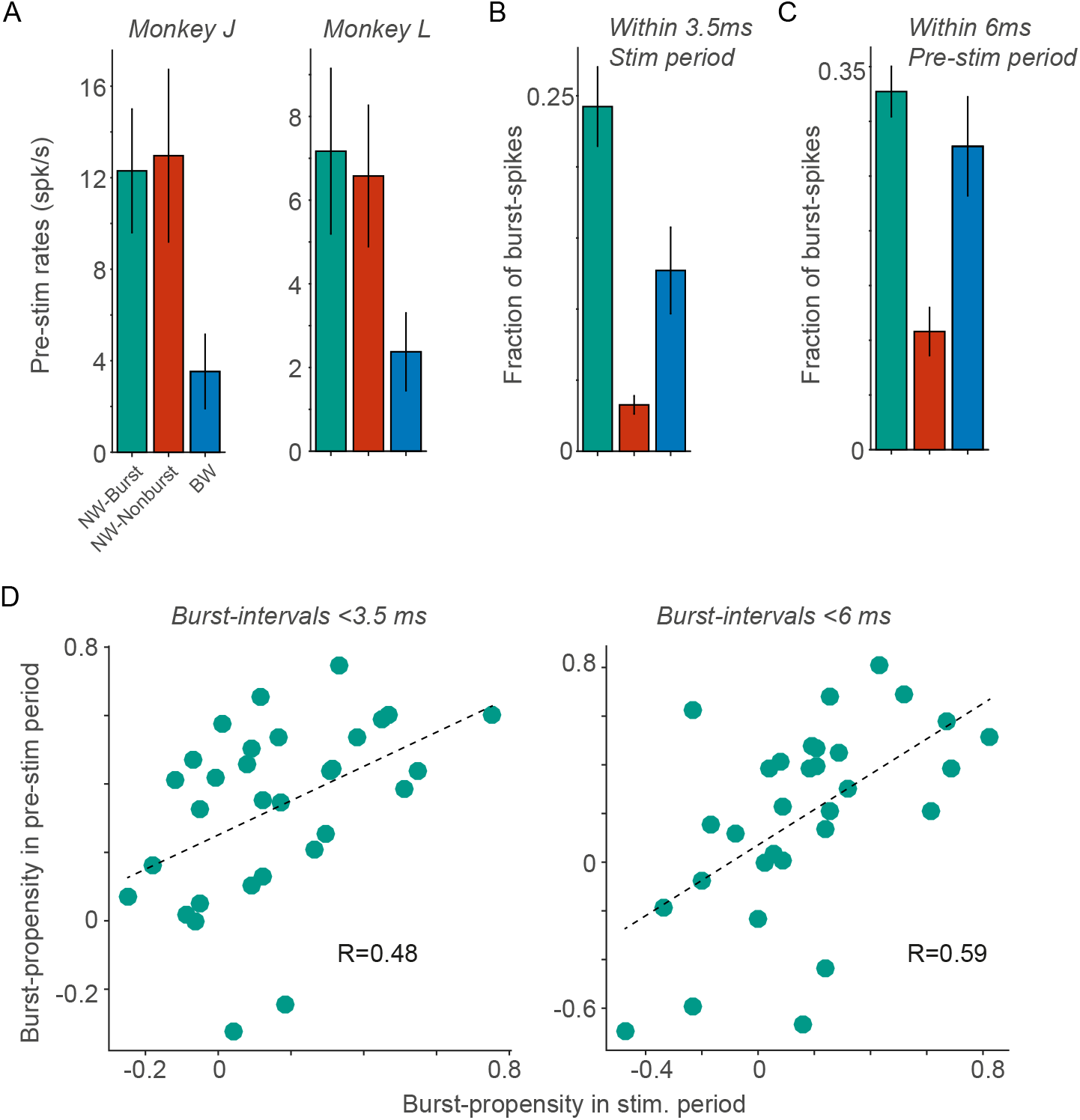
Neuron types differed in discharge rates and burst-propensity. The burst-propensity of NW–Burst neurons in the pre-stimulus period was high and correlated with the burst-propensity in the stimulus period. Relates to Figure 3 of the main text. (A) Similar to Figure 3B, now for monkeys J and L separately. (B) Similar to Figure 3A, but now considering only burst-spikes within 3.5ms. NW–Burst neurons fired more burst-spikes than the other neuron types (BW: P<0.005; NW–Nonburst: P<0.001, permutation test). BW neurons fired more burst-spikes than NW–Nonburst neurons (P<0.005, permutation test). (C) Similar to Figure 3A, but now for the pre-stimulus period. (D) Correlation of burst-propensity between the pre-stimulus and the stimulus period for NW–Burst neurons. Left: burstiness defined based on intervals < 3.5s. Right: burstiness defined based on intervals < 6ms. There was no significant difference in burst-propensity between the pre-stimulus and stimulus period (*P* > 0.05, paired T-test).

**Figure S5:**
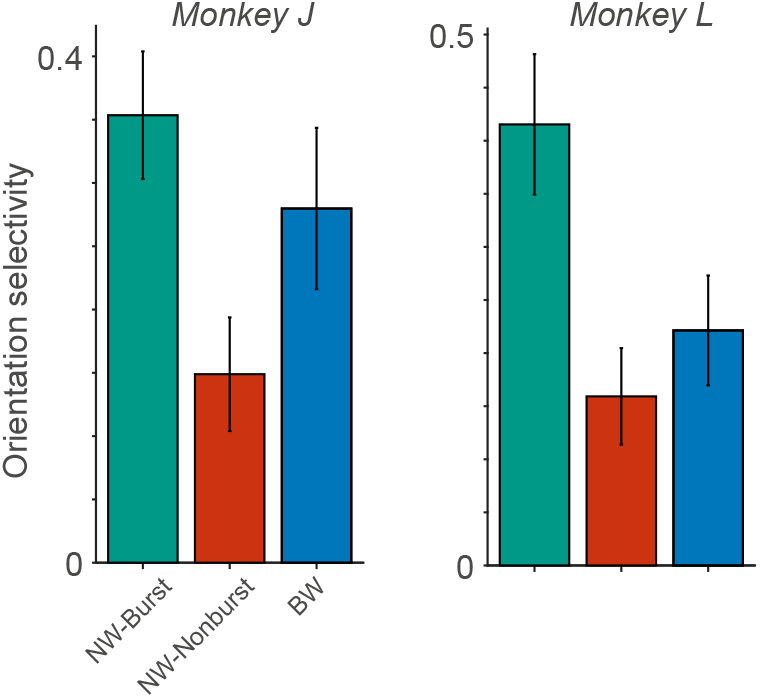
Analysis of OSI for two monkeys separately. Relates to Figure 4 of the main text. Similar to Figure 4A, but now for the two monkeys separately.

**Figure S6:**
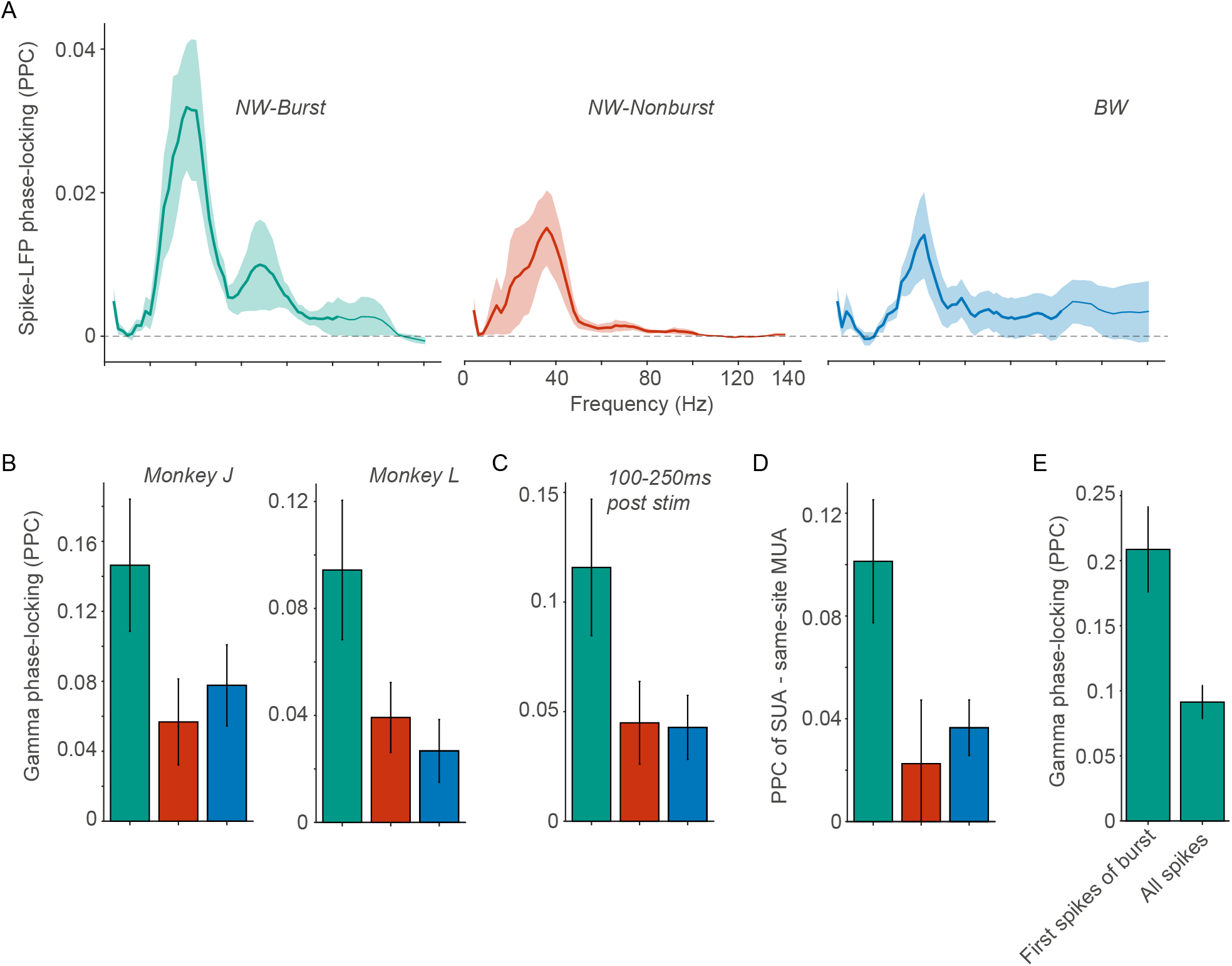
NW–Burst neurons were more gamma phase-locked than NW–Nonburst and BW neurons, relates to Figure 6 in main text. (A) Similar to Figure 6A, but now for monkey L. (B) Similar to Figure 6B, but now separately for monkeys J and L. (C) Similar to Figure 4B, but now for the 100-250ms period. Note that panels C-E combine both monkeys. (D) Difference between gamma PPC for the single unit and the corresponding same-site MUA site. The same-site MUA was defined as the set of all spikes from an electrode after exclusion of the single unit. NW–Burst neurons were more gamma phase-locked than the same-site MUAs were. (E) Comparison of spike-LFP gamma phase-locking (PPC) between first spikes of the burst and all the spikes, for NW–Burst neurons. First spikes of a burst were defined as spikes that were preceded by a pause of at least 10ms, and followed by a spike within 3.5ms. First spikes of burst were significantly more gamma phase-locked than all spikes together (*P* < 0.001, T-Test).

